# Fully-primed slowly-recovering vesicles mediate presynaptic LTP at neocortical neurons

**DOI:** 10.1101/2023.04.11.535831

**Authors:** Iron Weichard, Holger Taschenberger, Felix Gsell, Grit Bornschein, Andreas Ritzau-Jost, Hartmut Schmidt, Robert J. Kittel, Jens Eilers, Erwin Neher, Stefan Hallermann, Jana Nerlich

## Abstract

Pre- and postsynaptic forms of long-term potentiation (LTP) are candidate synaptic mechanisms underlying learning and memory. At layer 5 pyramidal neurons LTP increases the initial synaptic strength but also short-term depression during high-frequency transmission. This classical form of presynaptic LTP has been referred to as redistribution of synaptic efficacy. However, the underlying mechanisms remain unclear. We therefore performed whole-cell recordings from layer 5 pyramidal neurons in acute cortical slices of rats and analyzed presynaptic function before and after LTP induction by paired pre- and postsynaptic neuronal activity. LTP was successfully induced in about half of the synaptic connections tested and resulted in increased synaptic depression during high-frequency transmission and a decelerated recovery from depression due to an increased occurrence of a slow recovery component. Analysis with a recently established sequential two-step vesicle priming model indicates an increase in the abundance of fully-primed and slowly-recovering vesicles. A systematic analysis of short-term plasticity and synapse-to-synapse variability of synaptic strength at various types of synapses revealed that stronger synapses generally recover more slowly from synaptic depression. Finally, pharmacological stimulation of the cyclic adenosine monophosphate (cAMP) and diacylglycerol (DAG) signaling pathways, which are both known to promote synaptic vesicle priming mimicked electrically-induced LTP and slowed the recovery from depression. Our data thus demonstrate that LTP at layer 5 pyramidal neurons increases synaptic strength primarily by enlarging a subpool of fully-primed slowly-recovering vesicles.

## Significance statement

Despite intense investigation, the mechanisms of presynaptic LTP remain poorly understood. This is in part due to an incomplete knowledge of presynaptic function itself. The molecular process of generating fusion competent synaptic vesicles is referred to as priming. Many of the proteins involved in vesicle priming have long been identified but only recent studies revealed significant heterogeneity among primed synaptic vesicles (such as normally and superprimed or loosely and tightly docked vesicles). Here we aimed to analyze presynaptic LTP in light of these recent findings. We show that heterogeneities in vesicle priming must be considered to provide a mechanistic understanding of presynaptic LTP.

## Main Text

### Introduction

Experience-driven strengthening of synapses is the best-understood candidate mechanism mediating learning and memory (1). Long-lasting changes in synaptic strength can be caused by changes in presynaptic neurotransmitter release and/or postsynaptic receptor function. Because presynaptic mechanisms of LTP are less well understood and uncovering their molecular mechanisms and pathways is of fundamental importance for understanding physiological and pathophysiological processes during learning and memory formation (2), presynaptic LTP has been studied intensely in recent years in several brain regions, including hippocampus, neocortex, amygdala, thalamus, and cerebellum (3–5).

For presynaptic LTP observed at large hippocampal mossy fiber boutons it is generally assumed that induction and expression occur presynaptically (3). However, the underlying mechanisms remain incompletely understood because general aspects of synaptic vesicle priming and fusion and their relation to the regulation of presynaptic function during LTP are still unresolved. Even more limited is our understanding of presynaptic LTP at small conventional boutons mostly exhibiting postsynaptic N-methyl-D-aspartate receptor (NMDAR)-dependent induction of LTP. While the relevance of presynaptic LTP at conventional boutons is still debated (6), it is widely accepted that presynaptic LTP can occur at small conventional boutons exhibiting NMDAR-dependent LTP induction (7, 8) and that it is important for memory formation (2, 3).

A prototypical example of presynaptic LTP at conventional excitatory boutons has been described at synapses onto neocortical layer 5 pyramidal cells (9–11), where LTP is induced at the postsynaptic site via activation of NMDAR (12–15). At these synapses an increase in synaptic strength is accompanied by an increase in short-term depression during high-frequency transmission, referred to as redistribution of the synaptic efficacy (9, 16), i.e., synaptic efficacy increases for the first but decreases for subsequent stimuli. An increased probability of release of fusion-competent vesicles (17), also referred to as vesicular release probability (p_vr_), represents one possible mechanism for such redistribution as it augments early release but attenuates late release due to increased exhaustion of the vesicle pool (9, 17).

Recent advances in our molecular understanding of presynaptic function suggest that vesicle priming is a multi-step process resulting in different degrees of priming (18–25). Consistent with this notion, high-resolution structural analyses revealed heterogeneous degrees of tethering and docking of vesicles (26–28) and vesicles residing in such structurally distinct priming states have recently been referred to as loosely and tightly docked vesicles (24) or replacement and docked vesicles (29). Furthermore, functional analyses provided evidence for heterogeneous p_vr_ among release-ready vesicles, giving rise to different vesicle subpools referred to as primed and pre-primed (30), normally primed and superprimed (31, 32), loosely and tightly docked (33), and replacement and docked vesicles (29). Assuming that these molecular, structural, and functional definitions reflect a common concept that release machinery assembly represent a sequential molecular multi-step process giving rise to structurally and functionally distinct vesicle priming states (27), we use the umbrella term fully-primed for vesicles in the higher priming state throughout the manuscript. The main aim of this study was to analyze the mechanisms of presynaptic LTP in the context of the recent indications for heterogeneous vesicle priming. Our results show that fully-primed vesicles recover intrinsically slower from synaptic depression and their increased abundance primarily mediates presynaptic LTP at layer 5 pyramidal cell input synapses.

## RESULTS

### LTP of excitatory synapses onto neocortical layer 5 pyramidal neurons

To study long-term changes in synaptic strength at excitatory synapses onto neocortical layer 5 pyramidal cells, EPSCs were evoked by extracellular stimulation of excitatory inputs (Fig. 1A). LTP was induced electrically by pairing extracellular axonal stimulation with postsynaptic supra-threshold depolarization as previously described (9, 10) (Fig. 1B and Methods). To obtain estimates for release probability and vesicle pool size, paired-pulses and high-frequency stimulations were repeatedly applied before and after LTP induction (Fig. 1B). LTP induced a significant increase of the initial EPSC amplitudes in 21 out of 39 cells tested (referred to as responder cells; Fig. 1C). In responder cells, the median of EPSC amplitudes increased from 132 pA [124 pA, 151 pA] to 202 pA [164 pA, 251 pA] (Fig. 1E), corresponding to a relative increase by 47% [23%, 77%] (median value [first quartile, third quartile]; Fig.1F; p<0.001; n=21). In the remaining 18 cells, stimulation with the LTP induction protocol failed to increase initial EPSC amplitudes (p>0.05 for each cell, referred to as non-responder cells). Consistently, the average EPSC amplitude across all non-responder cells was unaltered after stimulation with the LTP induction protocol (Fig. 1E, p=0.27). Importantly, in the combined dataset of responder and non-responder cells, the EPSC amplitude significantly increased after stimulation with the LTP induction protocol (p<0.001; n=39; Fig. 1E), excluding the possibility that the increase in EPSC amplitudes in the responder cells was due to selecting cells with randomly increasing EPSC amplitudes. To ascertain stability of recordings, stimulation with the LTP induction protocol was omitted in some cells. These cells showed stable EPSC amplitudes throughout the recording period (control cells, p=0.46, n=8, Fig. 1E and F). Taken together, these data indicate that the pairing of pre- and postsynaptic activity induces robust LTP in about half of the inputs onto layer 5 pyramidal neurons.

**Fig. 1.**
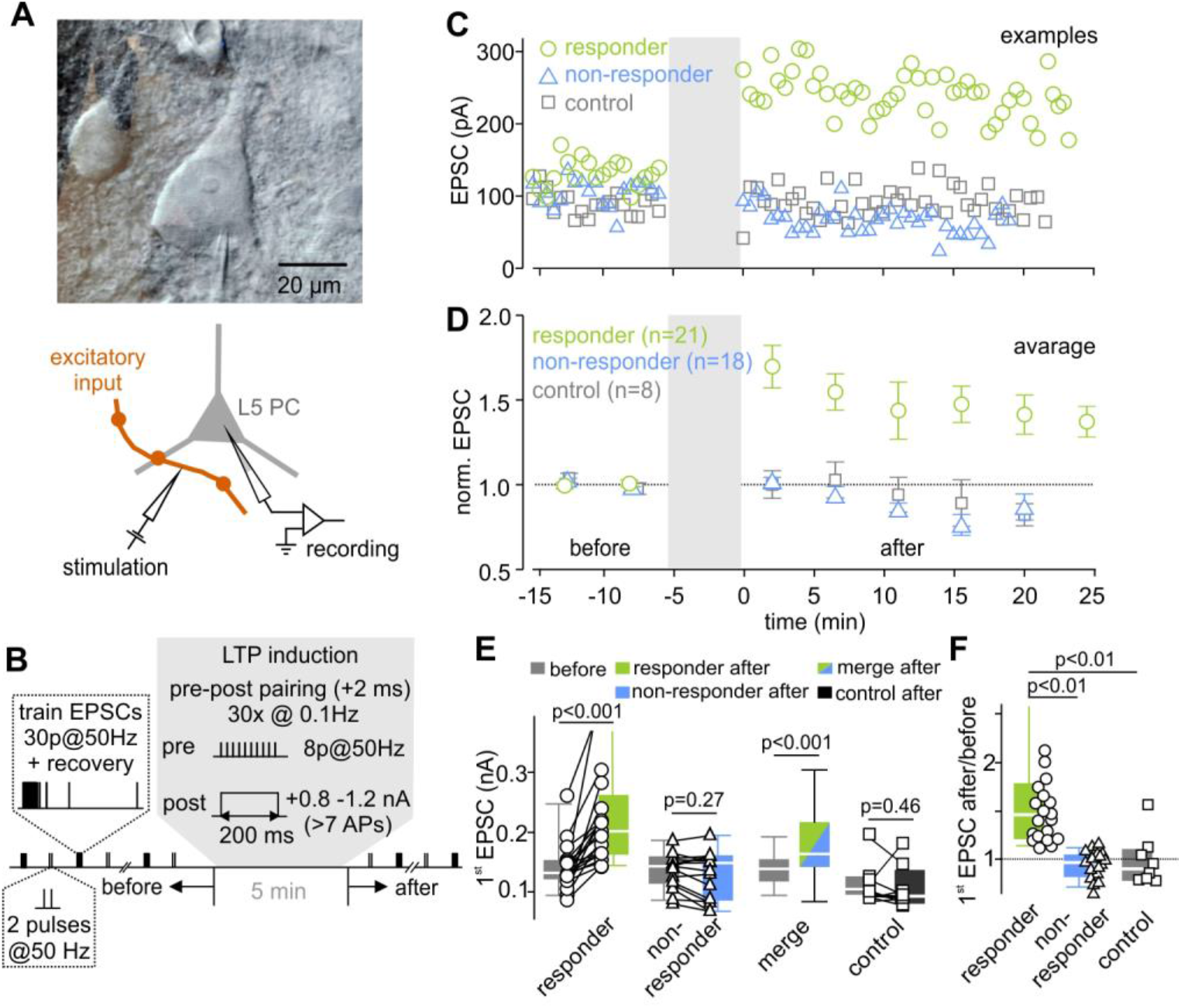
LTP of excitatory synapses onto neocortical layer 5 pyramidal neurons. (A) Infrared DIC image of a patch-clamped layer 5 pyramidal cell (top) and schematic illustration of the recording and stimulation configuration (bottom). (B) Illustration of the stimulation protocol used to assess short-term plasticity changes associated with LTP induction in layer 5 pyramidal neurons. Inputs were repeatedly stimulated extracellularly with 50 Hz trains consisting of 30 stimuli followed by single stimuli delivered at various time intervals after a conditioning 50 Hz train to probe recovery from depression. In addition, paired-pulses with 20 ms inter-stimulus interval were applied after 20 s. This stimulation pattern (train + paired-pulses) was repeated every 45 s for 10 min. To induce LTP, pre- and postsynaptic activities were paired while recording in current-clamp configuration. (C) Three example experiments of LTP responder (green), non-responder (blue), and control cells (grey, receiving no LTP induction). (D) Normalized mean EPSC amplitudes of responder, non-responder and control cells before and after LTP induction. (E) Individual and median EPSC amplitudes of responder and non-responder cells, of the merged dataset (responder and non-responder), and of a control group (no stimulation). (F) Box plot and individual data points of the change in EPSC amplitude (after / before) of responder, non-responder and control cells.

EPSCs of responder and non-responder layer 5 pyramidal cells had comparable amplitudes and kinetics before LTP induction (Fig. S7), suggesting the recruitment of the same number and type of inputs. Furthermore, short-term plasticity and recovery from synaptic depression (described in more details below) measured before LTP induction were similar in responder and non-responder cells (Fig. S7). In addition, in paired recordings of synaptically coupled layer 5 pyramidal cells (n=12 layer 5 pyramidal cell pairs), a significant increase in synaptic strength after LTP induction was observed in a subset of layer 5 pyramidal cell pairs. As for extracellular axonal stimulation, in paired recordings half of the layer 5 pyramidal cells showed an EPSP potentiation (n=6 of 12 tested; Fig. S3), whereas the other half of the cells did not show an increase of EPSP amplitudes following stimulation with the induction protocol. Thus, in about half of the layer 5 - layer 5 connections, LTP was successfully induced under our experimental conditions regardless of the mode of stimulation (Figs. 1 and S3). Similar observations were made previously at these synapses, where the same stimulus protocol exhibits LTP in only a fraction of the cells (10). Furthermore, the success rate of LTP induction depends on the stimulus intensity (11) and on the calcium signals in the postsynaptic dendrites (13).

### LTP changes short-term plasticity, increases the number of readily-releasable vesicles, and slows the recovery from synaptic depression

To assess presynaptic mechanisms of LTP, we evaluated changes in short-term plasticity before and after LTP induction by analyzing the EPSC amplitudes during and following high-frequency stimulation (30 stimuli at 50 Hz). To exclude postsynaptic contributions to short-term plasticity we recorded EPSCs in response to high-frequency stimulation in the presence of γ-D-glutamylglycine (γDGG, 2-3 mM), a low affinity AMPAR antagonist, attenuating AMPAR saturation and desensitization (34). γDGG reduced the median of the initial EPSC amplitude to 39% [0.27%, 0.41%] (n=6, Fig. S2C) while paired-pulse ratio (PPR), steady-state depression level and time course of recovery from synaptic depression (see below) were unaffected (Fig. S2B and C). These data argue strongly against a postsynaptic contribution and rather indicate that short-term plasticity at the stimulated synapses is mediated by presynaptic mechanisms, such as vesicle priming kinetics and pool depletion.

Provided that PPRs are mainly governed by vesicle depletion, changes in PPR may indicate changes in release probability (PPR ∝ (1-p_vr_); but see (33) that PPR and p_vr_ are not necessarily related). Following LTP induction, PPRs decreased by 10% [2.5%, 14%] from 0.85 to 0.77 in responder cells (Fig. 2C, p=0.004, n=21), but remained unchanged in non-responder cells (p=0.87, n=18). Furthermore, relative short-term depression increased after LTP induction in responder cells as indicated by their smaller normalized steady-state EPSCs amplitudes during high-frequency transmission (EPSC_steady-state_ / EPSC_1_). The normalized steady-state EPSC decreased by 20% [13%, 31%] from 0.26 [0.22, 0.35] to 0.21 [0.18, 0.26] (p≤0.001) in responder cells, but remained unchanged in non-responder cells (p=0.5) (Figs. 2E and S8). The observed changes in short-term plasticity suggest an increase in the p_vr_ during LTP.

**Fig. 2.**
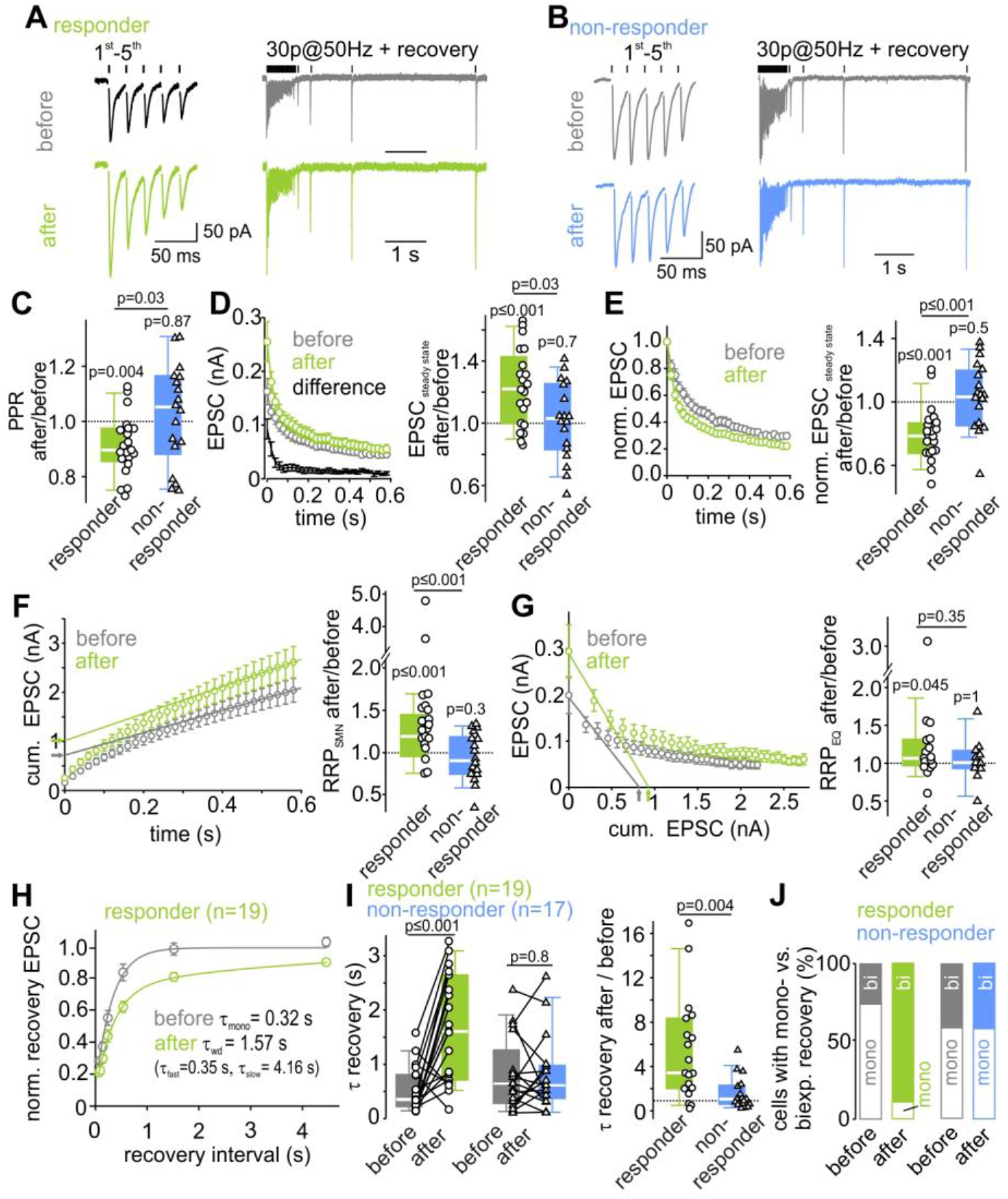
LTP changes short-term plasticity, increases the number of readily-releasable vesicles, and slows the recovery from synaptic depression. (A, B) Example average traces of the first 5 EPSCs (left) of a 50 Hz-EPSC train (right), followed by EPSCs elicited by single stimuli at different inter-stimulus intervals to probe recovery from depression, recorded before (top, grey) and after (bottom, green) LTP induction in a responder (A) and in a non-responder cells (B). (C) Individual and median relative changes in PPR after LTP induction in responder and non-responder cells. (D) Mean EPSC amplitudes during 50-Hz-trains were increased after LTP induction in responder cells (left). The difference curve indicates an increased synaptic strength throughout the train with a higher potentiation for the first EPSCs in responder cells. (Right) Relative changes of steady-state EPSC amplitudes after LTP induction in both responder and non-responder cells. (E) Mean normalized EPSC trains exhibit short-term depression, which increased in responder cells after LTP induction. (F) RRP size estimation by back-extrapolation of a linear fit to the last 5 amplitudes of the cumulative EPSC train plotted as a function of stimulus number (SNM method) shows an increase in responder cells (left). Relative changes of RRP size after LTP induction for responder and non-responder cells are shown in the right panel. (G) RRP size estimation by forward-extrapolation of a linear fit to the first 3 amplitudes of the EPSC train plotted as a function of cumulative previous release (EQ method) also shows an increase after LTP induction in responder cells. Relative changes of RRP size after LTP induction for responder and non-responder cells are shown in the right panel. (H) Mean normalized EPSCs during recovery from depression plotted as a function of recovery interval reveal a slower recovery from synaptic depression after LTP induction in responder cells. The time constants of recovery from depression were calculated from either mono- or bi-exponential fits to the mean recovery time course (lines). (I) Comparison of individual and median absolute (left) and summary plot of relative changes (right) of the weighted recovery time constant reveals a slowing of recovery from depression after LTP induction in responder but not in non-responder cells. (J) Proportion of cells (in %) with a mono- or bi-exponential recovery before and after LTP induction in responder and non-responder cells. Before LTP induction, a majority of the responder cells exhibits a mono-exponential recovery time course. After LTP induction, nearly all responder cells exhibit a bi-exponential recovery from depression.

Both the initial EPSCs and also the absolute steady-state EPSC of high-frequency EPSC trains were potentiated following LTP induction (cf. Fig. 2D). A sole redistribution of synaptic efficacy (i.e. an isolated p_vr_ increase) cannot account for the observed increase in steady-state EPSC. One possible reason could be changes in the readily releasable pool (RRP) of synaptic vesicles. To assess changes in the RRP we relied on established analyses of cumulative EPSC amplitudes (see Methods). Two approaches to estimate RRP size (35) indicated an increase between 7% and 30% in responder cells (7% [0%, 31%] with the EQ method, p=0.04, n=17, Fig. 2G; and 30% [15%, 54%] with the SMN method, p≤0.001, n=21, Fig. 2F). The RRP size was unchanged in non-responder cells (Fig. 2F and G and Fig. S8B and C; SMN method: p=0.3, EQ method p=0.99). Alternatively, the elevated steady-state amplitude can be explained by an increase in the quantal size. This is difficult to measure specifically for the potentiated synapse. We therefore used spontaneous EPSCs upon pharmacological potentiation of all synaptic inputs as an indirect surrogate. The observed increase of less than 6-12 % (see below; Fig. 5C) was smaller than increase in the RRP estimates (7-30%). These data suggest that the enlargement of the RRP contributes to LTP expression.

In addition, we observed changes in the recovery from depression (Fig. 2H). The recovery from short-term depression provides valuable information on the mechanisms of vesicle priming. Particularly, the number of exponential components and their relative contribution and time constant may reflect heterogeneity of vesicle priming and reveal details about the kinetics of vesicle priming (36). The recovery time course was therefore fit with either a mono- or bi-exponential function (for criteria see Methods) yielding either a single (τ_mono_) or two time constants (τ_fast_ and τ_slow_) with corresponding fractional amplitudes. In case of bi-exponential fits, a weighted time constant (τ_wd_) was calculated for comparison to single-exponential fits (cf. Methods). In responder cells, the median recovery time constant increased ∼5-fold from 0.35 s [0.22 s, 0.81 s] before to 1.61 s [0.74 s, 2.58 s] after LTP induction (Fig. 2I, p≤0.001, n=19). Before LTP induction, the EPSC recovery time course was best fit with a mono-exponential function in the majority of cells (Fig. 2J, τ_mono_=0.25 s [0.21 s, 0.38 s], n=14). After LTP induction, the recovery time course was best fit with a bi-exponential function in nearly all responder cells. In a majority of cells, an additional slow recovery component emerged after LTP induction (τ_slow_=4.39 s [1.98 s, 6.52 s], n=17), while τ_fast_ obtained from the bi-exponential recovery function was similar to τ_mono_ obtained from the mono-exponential recovery fit before LTP (τ_fast_=0.31 s [0.24 s, 0.43 s], n=17; τ_mono_ before vs. τ_fast_ after LTP p=0.46). In five cells the recovery was bi-exponential already before LTP. The τ_slow_ values were similar before and after LTP induction (unpaired Mann-Whitney-U-test p=0.39, n=5 vs. 17; paired Wilcoxon-test p=0.13, n=5 each; data not shown). Based on these observations, we conclude that the increase of τ_wd_ following LTP induction is caused by an increase in the relative contribution of the slow recovery component. The recovery from synaptic depression in non-responder cells was unaffected by the LTP induction protocol (Fig. 2I and J and Fig. S8). These findings are reminiscent of the previously observed additional slow component of recovery upon superpriming of vesicles with the GTP-binding protein Rab3 (31). In sum, the data show that recovery from synaptic depression becomes substantially slower after LTP induction by adding a second, slow recovery component, consistent with the hypothesis of an enlarged subpool of fully-primed slowly-recovering vesicles during LTP.

### Sequential and parallel models of vesicle priming can explain the short-term plasticity

To evaluate changes in vesicle priming before and after LTP induction more quantitatively, we tried to reproduce our experimental data in numerical simulations using two established models of vesicle priming and fusion. First, we tested a model in which release can originate from two parallel pathways represented by two types of fusion-competent vesicles states: high-p_vr_ vesicles, which recover slowly, and low-p_vr_ vesicles, which recover rapidly (Fig. S4A; referred to as ‘parallel model’ in the following). High-p_vr_ vesicles can be considered as being analogous to previously described “superprimed vesicles” (31, 32). In addition, a limited supply pool of vesicles is assumed (N_0_,(37)). Because high-p_vr_ vesicles intrinsically recover slowly, addition of high-p_vr_ vesicles after LTP induction can explain not only the observed increased EPSC amplitude and increased relative steady-state depression, but also the slower recovery of EPSC following high-frequency trains.

In addition, we tested a model that assumes two vesicle states (referred to as TS and LS for tightly and loosely docked states, respectively) of which only TS vesicles are fusion-competent (Fig. 3A; referred to as ‘sequential model’ in the following) (33). The recruitment of vesicles into LS and the transition from LS to TS is mediated by Ca^2+^-dependent rate constants. The LS to TS transition is slow at rest, but increases linearly with [Ca^2+^]_i_, such that each AP shifts a certain fraction of vesicles from LS to TS. The equilibrium between vesicles in states LS and TS at rest determines initial EPSC amplitudes. The balance between release of TS vesicles and their resupply determines the degree of the initial short-term depression including the PPR. A shift in the equilibrium between vesicles states at rest in favor of TS accounts for the observed increased EPSC amplitude and enhanced relative steady-state depression after LTP induction. The need to convert a higher proportion of vesicles from LS to TS (after depletion of TS vesicles) leads to a larger slow component of recovery and its overall slowdown.

**Fig. 3.**
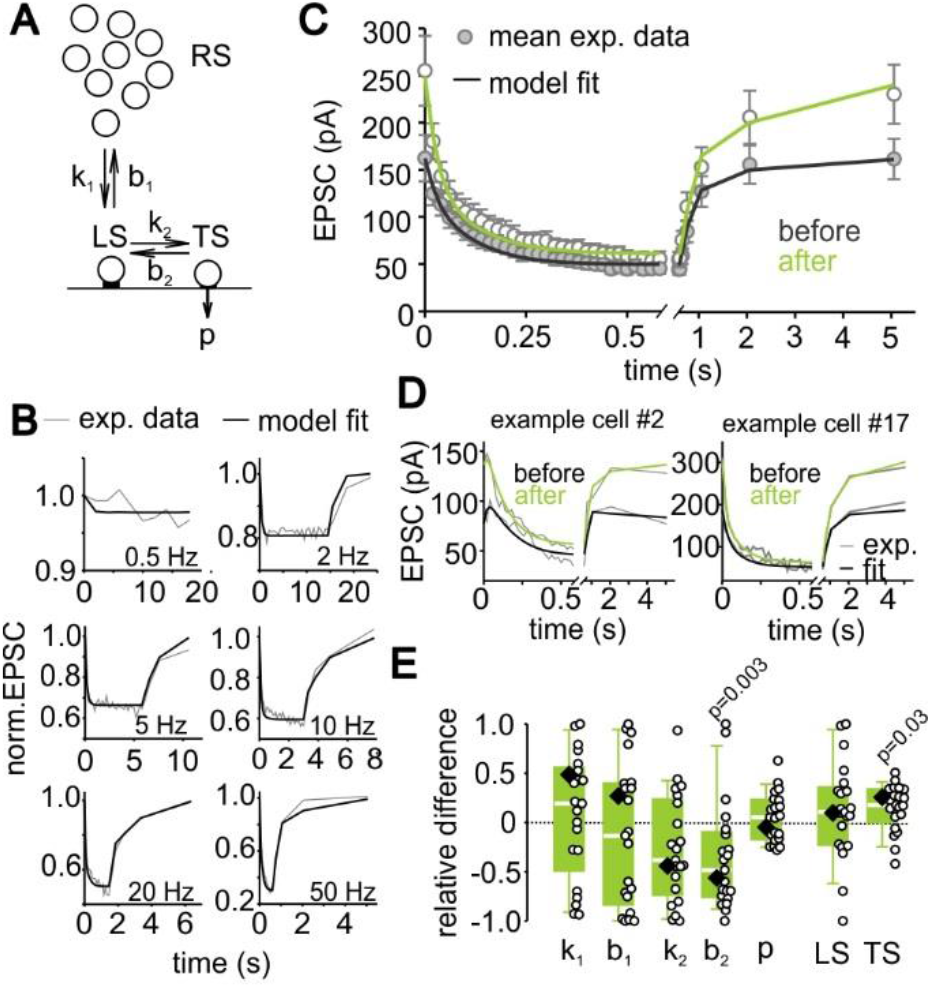
A sequential two-step priming model indicates an increased number of tightly docked vesicles following LTP induction. (A) Sequential model containing a loosely-docked vesicle subpool (LS), a fusion-competent tightly docked vesicle subpool (TS) and an infinite reserve pool (RS). (B) Model fit to average EPSC trains recorded in response to various stimulation frequencies. (C) Model fit to the mean 50-Hz EPSC train and the recovery EPSCs recorded in responder cells before and after LTP induction. (D) Model fits to individual 50-Hz EPSC trains and recovery EPSCs before and after LTP induction in two example cells. (E) LTP induced individual (circles) and median relative differences (eq. 7) of the predicted model parameters (diamond = parameter obtained from model fit to the average of all responder cells). P values were calculated by comparison of respective parameters before and after LTP induction by Wilcoxon signed rank tests.

We first compared the ability of the models to reproduce short-term depression during stimulus trains at frequencies ranging from 0.5 to 50 Hz and the subsequent recovery from depression. After optimization of the free parameters, both models reproduced short-term depression time courses well. The sequential model performed slightly better compared to the parallel model (Figs. 3B and S4B; χ^2^=15.97 and 19.34 for sequential and parallel model, respectively). We therefore show the results of the sequential and parallel model in main and supplementary figures, respectively. However, because both models are able to reproduce our experimental findings, we cannot unequivocally differentiate the biophysical plausibility of the models (38).

### Both models indicate an enlargement of a subpool of vesicles following LTP induction

Next, we optimized a subset of the five free parameters of both models (see Methods) to best reproduce short-term plasticity before and after LTP induction observed for the average response across all responder cells (Figs. 3C and S4C). In addition, we optimized the free parameters for each of the 21 connections individually (see Figs. 3D and S4D for two examples). We compared the best-fit parameters of the sequential model before and after LTP induction for each synaptic connection and plotted the relative difference or the ratio (Figs. 3E and S5; see eq. 7 in methods). The data indicate a significant reduction of b_2_, the rate constant of the backward transition TS → LS, which can be interpreted as a stabilization of the TS state after LTP induction and consequently a higher occupancy of the TS state at rest (see eq. 5 and 6 in methods; Fig. 3E). In analogy, the number of high-p_vr_ vesicles (N_2_) increased significantly when analyzing LTP induction-induced changes with the parallel model, (Fig. S4E). When non-responder cells were subjected to the same analysis no significant differences in model parameters before and after LTP induction were found (Fig. S8D-I). Thus, an unbiased parameter optimization in two mechanistically different models revealed a significant increase in the number of fully-primed slowly-recovering vesicles (TS vesicles in the sequential and N_2_ vesicles in the parallel model).

To corroborate that our results can be reproduced by assuming heterogeneous priming of subpools of vesicles, we used a linear unmixing-approach in which release during high-frequency trains observed in each of the 21 connections before and after LTP induction is described by a linear combination of components representing the distinct vesicle subpools of the models. We observed an exclusive increase in the TS component in the sequential and the N_2_ component in the parallel model upon inducing LTP (Fig. S6). These data therefore provide additional support for our hypothesis that LTP is mediated by an augmentation of the subpool of fully-primed slowly-recovering vesicles.

### Stronger synapses recover more slowly from depression

At the calyx of Held synapse, stronger synapses were shown to exhibit stronger paired-pulse depression (i.e. lower PPR), which was interpreted as a higher degree of superpriming (32) or else, as a higher proportion of TS vesicles (33). PPRs and initial EPSC sizes in layer 5 pyramidal cells obtained before LTP induction show a similar correlation between PPR and initial EPSC-amplitude (grey + black points in Fig. 4A; p=0.008). Strikingly, PPRs and initial EPSC sizes obtained after LTP induction follow the same relationship as seen before LTP induction extending the plot towards larger initial EPSCs. Thus, LTP “shifts synapses” along this relationship (cf. arrows in Fig. 4A and B and Fig. S3F for layer 5 pyramidal cell pairs). Since this correlation was previously associated with superpriming or a LS → TS shift, these data are consistent with fully-primed vesicles mediating LTP-induced enhancement of synaptic strength. In addition, we observed a strong correlation between the degree of relative steady-state depression and the kinetics of recovery in control data (before LTP induction; grey + black points in Fig. 4C, p<0.0001). Note, due to negative correlation of 1^st^ EPSC and relative steady-state EPSC amplitudes (data not shown; r_s_=-0.28, p 0.018, n=71), synapses with a low relative steady-state level have a high initial EPSC size and a slower recovery from depression. Again, the respective data obtained after LTP induction follow the same relationship between steady-state depression and τ of recovery, extending it towards slower recovery. In conclusion, these observations are consistent with our hypothesis that LTP induction primarily leads to a larger abundance of slowly-recovering vesicles (cf. arrow in Fig. 4C and D) or else shifting the LS ↔ TS equilibrium at rest towards TS.

**Fig. 4.**
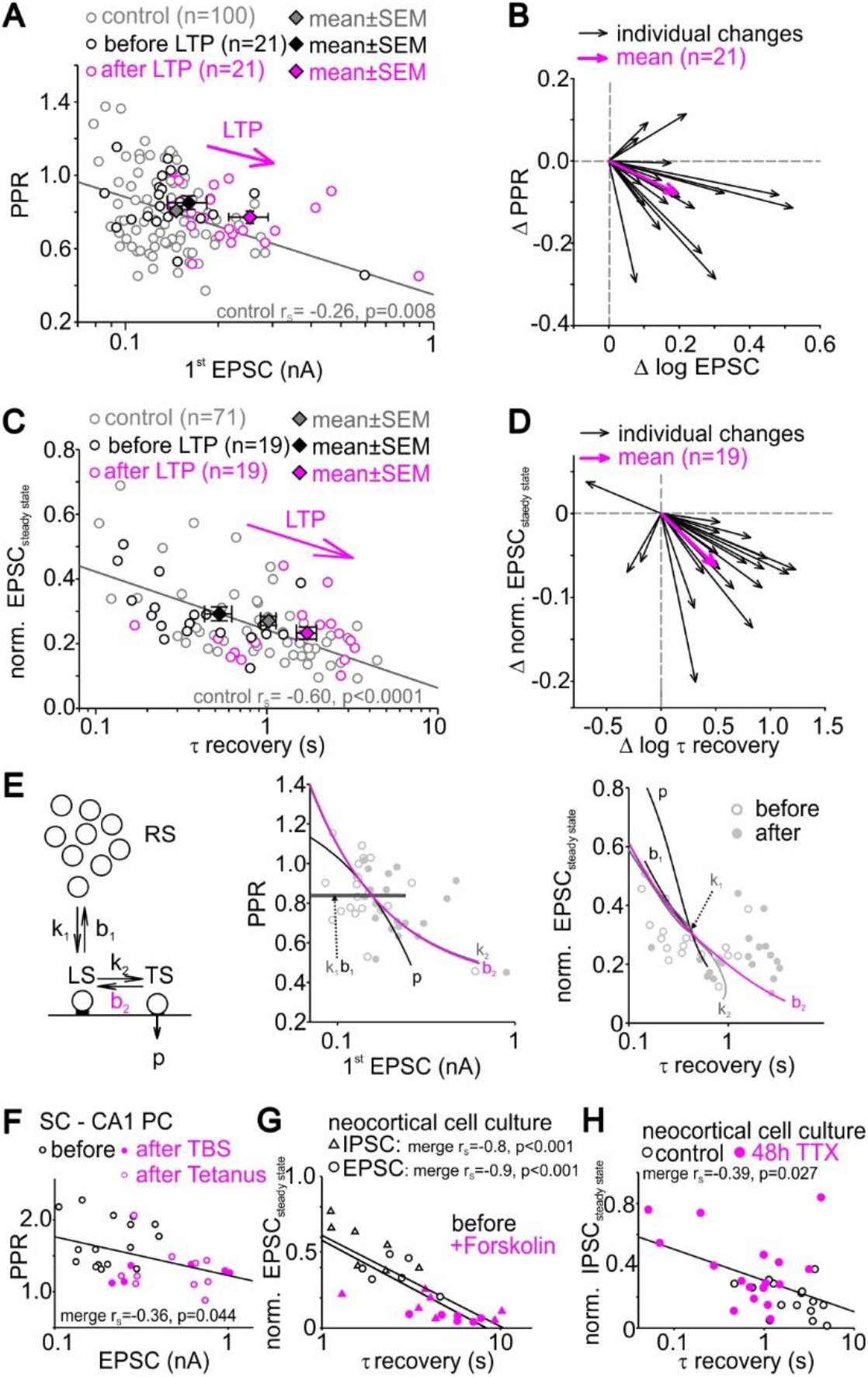
Stronger synapses recover more slowly from depression. (A) Scatter plot of PPR vs. EPSC size in layer 5 pyramidal cells before and after different after LTP induction. (B) Vector plot showing individual (black arrows) and average (magenta arrows) changes in EPSC size and PPR for responder cells (n=21). (C) Scatter plot of normalized steady-state EPSC vs. time constant of recovery from synaptic depression in layer 5 pyramidal cells before and after different LTP induction protocols. (D) Vector plot showing individual (black arrows) and average (magenta arrows) changes of normalized steady-state EPSC amplitudes and recovery from synaptic depression of responder cells (n=21). (E) Schematic of the sequential two-step priming model (left) and model predictions for the PPR vs. initial EPSC (middle) and normalized steady-state EPSC vs. recovery (right) relationships when systematically varying individual model parameters. (F) Scatter plot of PPR vs. EPSC size (cf. Fig 4A) before and after plasticity induction by theta burst (TBS)- and tetanic stimulation of Schaffer collateral to CA1 pyramidal cell synapses in acute brain slices. (G) Scatter plot of normalized steady-state PSC vs. time constant of recovery (cf. Fig. 4B) for high-frequency (20 pulses at 20 Hz) IPSC (triangles) and EPSC (circles) trains before and during forskolin application in synapses of cultured neocortical neurons. Postsynaptic currents shift towards stronger relative short-term depression and slower recovery from depression during forskolin application. (H) Scatter plot of normalized steady-state EPSC vs. time constant of depression (cf. Fig. 4B) for control high-frequency IPSC trains (20 pulses at 50 Hz) and after induction of homeostatic plasticity (48 h TTX) in synapses of cultured neocortical neurons. Note, the homeostatic plasticity-induced shift of IPSCs towards larger normalized steady-state IPSC and faster recovery from depression.

To explore possible mechanisms that may account for these correlations between initial EPSC and PPR as well as between normalize steady-state EPSCs and τ recovery, we systematically varied the free parameters of our two models during simulations. The two correlations were best predicted by varying b_2_ (LS←TS backward transition rate constant) in the sequential (Fig. 4E) and by varying N_2_ (size of the superprimed vesicle pool) in the parallel model (Fig. S4F and G) but less well by varying any of the other free parameters. These data indicate that the two analyzed correlations can only be explained by changing the abundance of TS vesicles in the sequential model (determined by b_2_) and by the number of superprimed vesicles (N_2_) in the parallel model, providing additional quantitative support to the notion that addition of fully-primed and slowly-recovering vesicles (TS or N_2_ vesicles) can explain both the variability among control synapses and their potentiation following LTP induction.

To test the synapse-type specificity of these correlations, we analyzed our unpublished data from other synapses. We observed similar correlations in hippocampal Schaffer collateral to CA1 pyramidal neuron synapses (Fig. 4F) and synapses between cultured neocortical neurons (Fig. 4G and H), not only with respect to the functional heterogeneity of synapses under control conditions (black symbols), but also with respect to plasticity induced changes by various means (magenta symbols). These data suggest that the abundance of fully-primed and slowly-recovering vesicles critically determines synapse-to-synapse variability of synaptic strength and short-term plasticity. Note, that the correlation between steady-state depression and τ recovery could not be assessed for hippocampal Schaffer collateral to CA1 pyramidal cell synapses in acute slices, which showed synaptic facilitation. Furthermore, the correlation between initial EPSC size and PPR could not be assessed for cultured neurons, which showed highly variable EPSC sizes within the unpaired control and treated cell group.

### Forskolin- and PDBu-induced potentiation slows recovery from depression

To further test our hypothesis of enhanced vesicle priming following LTP induction, we potentiated synapses by stimulating the cAMP and DAG pathway (Fig. 5A) via application of the adenlyate cyclase activator forskolin and the DAG analog PDBu. These two pharmacological manipulations have been shown to stimulate vesicle priming by modulation of active zone priming proteins including RIM1α and MUNC13, and thereby inducing an increase in synaptic strength (see Discussion).

**Fig. 5.**
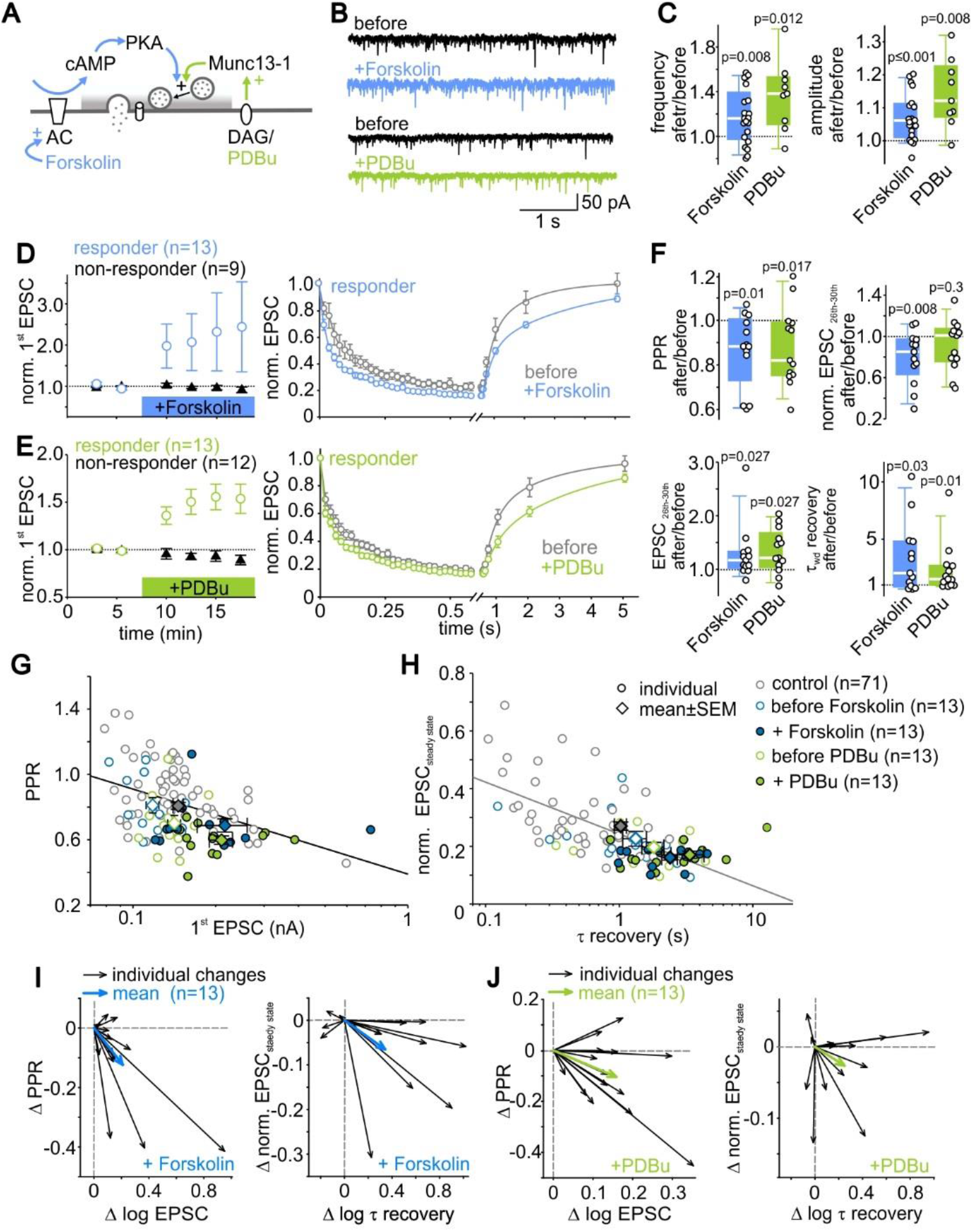
Forskolin- and PDBu-induced potentiation slows recovery from depression. (A) Schematic of forskolin- and PDBu-activated signaling pathways. (B) Example traces of spontaneous EPSCs (sEPSCs) before (black) and after (colored) forskolin or PDBu application. (C) Individual and median relative changes in sEPSC frequency (left) and amplitude (right) before and after forskolin or PDBu application. For sEPSC analysis, data from responder and non-responder cells were pooled. (D, E) Mean normalized initial EPSC amplitudes of responder and non-responder cells (left) and mean normalized 50-Hz EPSC trains and recovery EPSCs of responder cells (right) before and during forskolin (D) or PDBu (E) application. (F) Individual and mean relative changes in PPR, absolute and normalized steady-state EPSC amplitudes and recovery time constant in responder cells after forskolin or PDBu application. (G) Scatter plot of PPR vs. initial EPSC before and after induction of various forms of plasticity (control data same as in Fig. 4A). (H) Scatter plot of normalized steady-state EPSC amplitudes vs. recovery time constant before and after induction of various forms of plasticity (control data same as in Fig. 4C). (I) Vector plot showing individual (black arrows) and average (blue arrows) changes in EPSC size and PPR (left) and relative steady-state EPSC amplitudes and recovery from synaptic depression for forskolin responder cells (n=13). (J) Vector plot showing individual (black arrows) and average (blue arrows) changes in EPSC size and PPR (left) and relative steady-state EPSC amplitudes and recovery from synaptic depression for PDBu responder cells (n=13).

We first analyzed spontaneously occurring EPSCs (sEPSCs) before and during bath application of 40 µM forskolin or 1 µM PDBu (Fig. 5B). The median frequency of the sEPSCs increased by 16 % [0%, 36%] following forskolin application and by 38 % [12%, 52%] following PDBu application (Fig. 5C left, p=0.008, n=21; p=0.012, n=9; respectively), consistent with increased release because of higher release probability and/or increased number of docked vesicles (39). In addition, sEPSCs amplitudes increased after forskolin and PDBu treatment by a median percentage change of 6% [0%, 11%] (p<0.001, n=21) and 12% [1%, 22%] (p=0.008, n=9), respectively (Fig. 5C right). The median amplitude of the sEPSCs was 15.2 pA [13.4 pA, 16.1 pA] (n=30) under control conditions, which is comparable to previously reported amplitudes of spontaneously fusing vesicles (mEPSCs) (40). Therefore, a majority of sEPSCs might represent mEPSCs, which would indicate an increase of maximally 6-12% in quantal size. These data indicate that the pharmacologically induced potentiation is mainly mediated by presynaptic strengthening and that a contribution of postsynaptic potentiation is limited to 6-12%.

To evaluate the presynaptic mechanisms of pharmacologically induced EPSC potentiation, EPSCs (30 pulses at 50 Hz + recovery EPSCs) were repeatedly measured before and during bath application of forskolin or PDBu and analyzed with respect to changes in short-term synaptic plasticity and recovery from synaptic depression. As observed for LTP induced by electrical stimulation (cf. Figs. 1 and 2), in approximately half of the cells (responders), application of either forskolin or PDBu induced robust EPSC potentiation by 38% [27%, 56%] (n=13) and 37% [28%, 52%] (n=13), respectively (Fig. 5D and E), while in the remainder of the cells tested, EPSC amplitudes remained unchanged (non-responders). The fractions of cells exhibiting EPSC potentiation were similar for the different modes of LTP induction (electrical stimulation-induced LTP=54% forskolin-potentiation=59% PDBu-potentiation=52 %).

Strikingly, analysis of short-term plasticity during EPSC trains revealed decreased PPR, increased depression, and slower recovery from depression after application of either forskolin or PDBu (Fig. 5F), similar as observed for electrical stimulation-induced LTP (cf. Fig. 2). Consistently, corresponding shifts in the correlations were observed (Fig. 5G-J; cf. with Fig. 4A-D). These data indicate that irrespective of the mode of induction, potentiation of synaptic strength is accompanied by increased relative short-term depression and by a slowing of recovery after high-frequency transmission.

### Discussion

To investigate presynaptic mechanisms of LTP we analyzed short-term plasticity before and after LTP induction at excitatory synapses onto layer 5 pyramidal neurons. LTP induction by paired pre- and postsynaptic activity profoundly changed the short-term plasticity and slowed the recovery from depression. Synapse potentiation via stimulation of the cAMP or DAG pathway resulted in very similar changes. Fitting constrained short-term plasticity models to the data indicates that presynaptic LTP at layer 5 input synapses is mediated primarily by an increased abundance of fully-primed and slowly-recovering vesicles.

### Pre- versus postsynaptic LTP at neocortical synapses

While presynaptic LTP at large hippocampal mossy fiber synapses is well established (41, 42), involvement of presynaptic mechanisms in the early expression of LTP has been questioned altogether for small synapses that exhibit postsynaptic NMDAR-dependent LTP induction (6). The finding that postsynaptically “silent” synapses can be rapidly transformed into functional synapses by insertion of AMPARs during LTP induction argued in favor for a primarily postsynaptic site of LTP expression (43, 44). However, the observed changes in short-term plasticity cannot be explained by postsynaptic AMPA receptor insertion. Furthermore, AMPA receptor saturation and desensitization contributed little to the observed short-term plasticity (Fig. S2) and therefore cannot explain the altered short-term plasticity during LTP. Our data do not rule out a contribution of postsynaptic mechanisms to LTP expression at layer 5 synapses. Indeed, a small increase in the size of spontaneously occurring EPSCs was observed following chemical induction of LTP (Fig. 5C), which is likely of postsynaptic origin. However, our detailed quantitative and modelling analysis indicates that presynaptic mechanism account for a majority of the LTP-induced potentiation, consistent with previous studies showing a presynaptic component of LTP at small synapses in the CA1 region of the hippocampus (28, 45, 46) and the neocortex (10, 11).

### A sub-population of release-ready vesicles mediates LTP

In principle, presynaptic LTP can be mediated by changes in the number of release sites, the number of release-ready vesicles, and p_vr_. Even though Katz postulated the existence of release sites (N; (47)) already decades ago, estimating N experimentally is still complicated by the existence of release sites not occupied by a vesicle (48) and heterogeneities among release-ready vesicles due to superpriming (31, 32) or differences in the docking state (24). At layer 5 pyramidal cell input synapses, the increased short-term depression upon LTP induction was accounted for previously by increased p_vr_ (9). Our analysis indicates alternative or additional mechanisms involving an increase in the number of release-ready vesicles. Consistent with our results in the neocortex, an increased number of docked and active zone-associated vesicles has been observed at small CA1 hippocampal synapses following electrical and chemical LTP induction (28, 45). More docked vesicles have also been observed at the larger hippocampal mossy fiber boutons following chemical (42) and post-tetanic potentiation (49, 50) and at synapses of cultured hippocampal synapses following silencing-induced homeostatic plasticity (51).

In addition to changes in the number of release-ready vesicles, we found a slowing of the recovery from depression following LTP induction (Fig. 2H and I). The slower recovery was mediated by a selective increase in the amplitude of the slow component of recovery, resulting in more cells with bi-exponential time courses of recovery from depression following LTP induction (Fig. 2E). Such change in the recovery time course is difficult to explain by a sole increase in p_vr_ and/or the number of release-ready vesicles. Instead, our data argue for a heterogeneity among release-ready vesicles as shown e.g., at hippocampal synapses (31, 52), calyx of Held synapse (32, 53, 54), the cerebellar mossy fiber bouton (37, 55), and neuromuscular junctions of drosophila (56) and of cray fish (57). We extend this concept by showing that release-ready vesicles not only differ in their baseline function but also in their responsiveness to LTP. In particular, our data indicate that a sub-population of vesicles, which are fully-primed and slowly-recovering, is up-regulated following LTP induction. Our functional analysis is strikingly consistent with a recent electron microscopic study at CA1 hippocampal synapses reporting that the density of tightly docked vesicles increased following LTP induction, whereas the density of loosely docked vesicles remained unchanged (28). It is therefore tempting to speculate that the vesicles with shorter tethers represent fully-primed vesicles (58), which recover slowly from depression.

### Why is LTP slowing the recovery from depression?

Considering that LTP represents a strengthening of synapses, our discovery of a slower time course of recovery from synaptic depression in the potentiated synapses seems surprising. The slower recovery after LTP has implications for computational neuroscience. It can alter the temporal flow of information in a neuronal network beyond a mere strengthening of synaptic weights (16, 59, 60). We observed negative correlations between synaptic strength and the speed of recovery from synaptic depression in several types of synapses (Fig. 4), suggesting that the slower recovery represents biophysical limitations of the time required to build the mature release machinery of fully-primed vesicles under resting conditions.

A recent study inducing LTP at hippocampal mossy fiber boutons with optogenetic stimulation found an increased rate of vesicle replenishment (61). This apparent discrepancy with the here-observed slower recovery from depression can be understood as follows. In the study by Fukaya et al. (61), release was assessed with repetitive 20-ms depolarizations and capacitance measurements. This technique provides the absolute rate of vesicle replenishment per bouton. It is the product of the number of release-ready vesicles and the recruitment rate per vesicle and determines the steady-state EPSC amplitude during high-frequency transmission (cf. Box 1 in ref. (36)). Indeed, we observed an elevated steady-state EPSC amplitude in potentiated synapses (Fig. 2D) indicating that LTP increased the absolute rate of vesicle replenishment per bouton also at the here-studied neocortical synapses. Furthermore, the heterogeneities among release-ready vesicles and the activity-dependence of vesicle replenishment provide additional explanations for why recovery from depression is slower while the absolute rate of vesicle replenishment during activity is increased.

### Priming during electrical stimulation-induced LTP and chemical potentiation

Our data favor a mechanism based on an increase in the priming states of vesicles underlying electrical stimulation-induced LTP. To test our hypothesis of enhanced vesicle priming, we used pharmacological tools, which can activate active zone proteins known to facilitate vesicle priming. Munc13 is a key priming factor of synaptic vesicles exhibiting several regulatory domains including the C1 domain which binds DAG and can be activated by DAG analogues, such as the phorbol ester PDBu (62–65). Furthermore, the C2A domain of Munc13 can be activated through its interaction with Rim1α (66–69) and Rim1α in turn can be activated by the protein kinase A/cAMP pathway (70–72), but see (73). We therefore focused on the pharmacological activation of the DAG- and cAMP-pathway by PDBu and forskolin, respectively. We found striking similarities in the changes in short-term plasticity upon electrical stimulation-induced LTP and upon pharmacological stimulation of the cAMP or DAG pathways. Particularly, the increased appearance of a slow component in the time course of recovery occurring upon both electrical stimulation-induced LTP and pharmacological boosting of vesicle priming strongly supports our conclusion that LTP is mediated by enhanced vesicle priming.

The presynaptic molecular mechanisms operating during electrical stimulation-induced LTP are still not fully understood. Particularly, the time course and magnitude of cAMP and DAG elevation required for the expression of LTP are unknown and may differ between synapses. Indeed, comparison of electrical stimulation-induced and forskolin-induced LTP revealed mechanistic differences regarding the involvement of Rab3 (74), Rim1α (71), PKA (75), and Ca^2+^ channels (76, 77). However, we established a number of similarities between electrical stimulation-induced LTP and pharmacological synaptic potentiation, which may indicate a convergence of down-stream pathways eventually activating Munc13.

### Parallel versus sequential mechanisms of vesicle priming

Although the kinetic schemes of the two previously established models of vesicle priming (Figs. 3A and S4A) appear to be quite similar, there are fundamental mechanistic differences. The sequential model (33) assumes a constant number of release sites, which can be empty or occupied by vesicles in one of two states (LS or TS). Only TS vesicles are fusion-competent and the mean p_vr_ is considered identical among synapses. Transitions between states are reversible, Ca^2+^-dependent and may be quite fast. Variations in both synaptic strength and plasticity among synapses and also upon LTP induction are explained by different resting occupancies of LS and TS states, which are in a dynamic equilibrium with each other and the empty state. The parallel model (37), in contrast, assumes two types of release sites (N_1_ and N_2_), which vary in number among synapses and may change upon induction of LTP. The two types of release sites have distinct p_vr_ and priming kinetics, they are fully occupied at rest, and recruitment of vesicles to sites is Ca^2+^-independent. Both models reproduce short-term plasticity at layer 5 pyramidal cell synapses across a large range of frequencies almost equally well (Figs. 3B and S4B). The parallel model has the advantage of being intuitively tangible. The sequential model has the advantage of incorporating recent experimental evidence regarding reversibility of priming and incomplete occupancy of release sites (33, 78, 79), Ca^2+^-dependence of vesicle recruitment (78, 80, 81), and varying degrees of tight apposition between vesicle and active zone membrane of morphologically docked vesicles (26, 27). Accordingly, both models predict different mechanisms underlying LTP. For example, the parallel model interprets LTP induction as an increased number of N_2_ sites, resulting in an increase in the average p_vr_ of the combined N_1_ and N_2_ pools (the proportion of N_2_ vesicles increases, while the p_vr_ value remains unchanged for both N_1_ and N_2_ vesicles).

Within the parallel-model framework, the recently described increased release probability during LTP at hippocampal mossy fiber boutons (61) is consistent with our findings because the used presynaptic capacitance measurements provide the increased average p_vr_ of N_1_ and N_2_ vesicles. The sequential model, in which only TS vesicles are fusion-competent, can reproduce the LTP-induced changes in synaptic strength and plasticity data without requiring a change in p_vr_. However, both models are consistent with the existence of release sites with high p_vr_ (40) and rapid vesicle supply-pools (82), as described at mature layer 5 pyramidal cell input synapses. Further studies are needed to differentiate unequivocally between the two models. While the parallel model requires mechanisms for regulating the number of release sites, the sequential model postulates changes in the priming state of docked vesicles, for which the regulatory domains of Munc13 offer several possibilities (24, 65). Nevertheless, it is reassuring that two models with profoundly different underlying mechanisms both support our main conclusion that fully-primed and slowly-recovering vesicles mediate LTP.

In summary, our study combined recent evidence for multi-step processes of vesicle priming with detailed quantitative dissection of the mechanisms of presynaptic LTP. We show that an increase of a sub-population of vesicles mediates LTP. These vesicles exhibit a fully matured priming state, a slow recovery from depression, and presumably a tight docking state with short vesicle tethering filaments (28).

## METHODS

### Animals

Animals were handled in accordance with European (EU Directive 2010/63/EU, Annex IV for animal experiments), national and Leipzig University guidelines. All experiments were approved in advance by the federal Saxonian Animal Welfare Committee (T29/19). Patch-clamp recordings were made from neocortical layer 5 pyramidal cells and hippocampal CA1 pyramidal cells in in acute brain slices obtained from Sprague Dawley (SPRD) rats of either sex at postnatal (P) day P13-P21.

### Slice preparation

Animals were anesthetized by inhalation of isoflurane (Baxter Deerfield, IL) and killed rapidly by decapitation. Neocortical coronal slices (400 µm) containing the primary somatic sensory (S1) region and hippocampal slices (300-400 µm, as described previously (83)) were cut using a Leica VT1200 microtome (Leica Microsystems, Wetzlar, Germany) in ice cold (3-4 °C) dissection solution containing (in mM): 92 NMDG, 2.5 KCl, 0.5 CaCl_2_,10 MgCl_2_, 1.2 NaH_2_PO_4_, 30 NaHCO_3_, 25 glucose, 3 sodium pyruvate, 5 N-acetylcysteine, 5 sodium ascorbate, 20 HEPES, 1 kynurenic acid. Slices were incubated in the dissection solution for 5 min at 35°C. To reduce synaptic activity, slices were subsequently stored for at least 30 min in an intermediate storage solution at room temperature (RT) containing (in mM): 92 NaCl, 2.5 KCl, 2 CaCl, 1 MgCl_2_, 1.2 NaH_2_PO_4_, 30 NaHCO_3_, 25 glucose, 3 sodium pyruvate, 5 N-acetylcysteine, 5 sodium ascorbate, 20 HEPES and thereafter incubated in standard recording ACSF for at least 30 min before recordings containing (in mM): 125 NaCl, 2.5 KCl, 2 CaCl_2_, 1 MgCl_2_, 1.25 NaH_2_PO_4_, 25 NaHCO_3_, 25 glucose. This approach of using NMDG solutions was performed as previously described (84). During experiments, slices were continuously perfused with standard recording ACSF. All solutions were continuously equilibrated with 5% CO_2_ and 95% O_2_ to pH 7.4. Recordings were performed at near physiological temperature (33-36 °C).

### Cell culture

Neocortical neuronal cultures were prepared as described previously (85). In short, mice were decapitated and cerebral cortices were removed, dissected in ice-cold Hank’s balanced saline solution, and cut into 5–10 pieces per hemisphere. Brain tissue was digested by Trypsin (5 mg, dissolved in digestion solution; for composition see below) for 5 min at 37 °C. Trypsin digestion was stopped by ice-cold MEM growth medium (composition see below), and cells were mechanically dissociated using narrow-bore Pasteur pipettes in DNAse-supplemented MEM growth medium (10 mg/mL). Following two min of segregation by gravity, the supernatant was centrifuged and the resulting pellet was resuspended in MEM (Sigma) growth medium for plating onto Matrigel-coated (Corning®) 14 mm coverslips (50.000 vital cells/coverslip). Forty-eight hours after plating, the medium was fully replaced by MEM growth medium containing 4 µM Ara-C to limit glial growth, which was again fully replaced by fresh MEM growth medium 96 hours after plating. Cells were incubated at 37 °C, 93% humidity, and room air plus 5% CO_2_ until use. Digestion solution contained the following (in mM): 137 NaCl, 5 KCl, 7 Na_2_HPO_4_, 25 HEPES, pH adjusted to 7.2 by NaOH. MEM growth medium was prepared from 1l MEM (Earle’s salts + L-Glutamine) supplemented with 5 g Glucose, 0.2 g NaHCO_3_, 0.1 g bovine holo-Transferrin, 0.025 g bovine insulin, 50 mL FBS and 10 mL B-27.

### Electrophysiology

Postsynaptic whole-cell recordings were obtained from visually identified layer 5 pyramidal neurons in the rat somatosensory cortex S1 region, from pyramidal cells in the CA1 region of the hippocampus in acute brain slices, and from cultured neocortical neurons using an EPC 10 USB amplifier (HEKA Elektronik). Pipette solution for voltage-clamp recordings in acute brain slices contained (in mM): 130 KMeSO_3_, 10 KCl, 10 HEPES, 0.1 EGTA, 3 Mg-ATP, 0.3 Na-GTP, 5 Na-phosphocreatin, pH adjusted to 7.31 by KOH and osmolarity to 294 mOsm by sucrose. Pipette solution for voltage-clamp recordings of EPSCs in cultured neurons contained (in mM): 150 K-Gluconate, 10 K-HEPES, 3 Mg-ATP, 0.3 Na-GTP, 0.05 EGTA, 10 NaCl, 3 QX 314-Cl, pH adjusted to 7.35 by KOH, osmolarity 295 mOsm. Pipette solution for voltage-clamp recordings of IPSCs in cultured neurons contained (in mM): 40 CsCl, 10 HEPES, 90 K-gluconate, 0.05 EGTA, 10 Na-phosphocreatin, 1.8 NaCl, 2 MgATP, 0.4 Na2-GTP, 1.7 MgCl_2_, 3.5 KCl, 3 QX314-Cl, pH adjusted to 7.35 by CsOH, osmolarity 297 mOsm.

Recordings in acute brain slices were obtained using artificial cerebrospinal fluid (ACSF) solution as described above. In a subset of experiments, ACSF was supplemented with the low-affinity AMPAR antagonist γDGG (2-3 mM) to attenuate postsynaptic saturation and desensitization. Recordings in cultured neurons were obtained using an external solution containing (in mM): 150 NaCl, 4 KCl, 10 HEPES, 10 glucose, 1.1 MgCl_2_, 1.1 CaCl_2_, pH adjusted to 7.4 by NaOH at 35 °C, osmolarity 305 mOsm. For all experiments, external solution was supplemented with blockers of NMDARs (20 µM APV) and GABA_B_Rs (3 µM CGP-55845). For recordings of AMPAR-mediated EPSCs, external solution was supplemented with the GABA_A_R antagonist SR95531 (10 µM) and the low-affinity GluR antagonist kynurenic acid (1 mM). For recordings of GABA_A_R-mediated IPSCs, the external solution was supplemented with the AMPAR antagonist NBQX (20 µM) and the low-affinity GABA_A_R antagonist TPMPA (0.3 mM).

Pipettes were pulled from borosilicate glass with open-tip resistance between 3 to 5 MΩ. For recordings in acute brain slices, the series resistance (R_s_) was monitored and R_s_ compensation was dynamically adjusted every 2 min to yield a remaining uncompensated R_s_ of 10 MΩ (mean initial R_s_ values without compensation was 11.9 ± 2.9 MΩ, n=54). Neurons were held in voltage-clamp mode at a holding potential of −70 mV during pre- and post-LTP induction recording periods. Voltage-clamp recordings were sampled at 20-100 kHz and low-pass filtered at 3.9 kHz.

Spontaneous EPSCs (sEPSCs, Fig. 5B and C) were measured without stimulation of excitatory inputs and in the absence of TTX. sEPSCs thus possibly represent a mixture of postsynaptic responses generated by spontaneous fusion of single vesicles (miniature EPSCs) and by vesicle release elicited by spontaneously occurring presynaptic action potentials.

### Biocytin labeling

For morphological identification, a subset of layer 5 pyramidal cells was filled with biocytin (2 mg/ml) during recordings and postfixed in 4 % PFA at 4 °C overnight. Subsequently, slices were washed repetitively in TBS and TBS + 0.3% Triton X-100 and incubated with Cy2-conjugated streptavidin (5 µg/ml, Jackson Immunoresearch Lab.) for 2.5 hours at RT. After several washes in TBS + 0.3% Triton X-100, TBS and distilled water, slices were mounted in Aqua-Poly/Mount (Polyscience) and inspected by confocal laser scanning microscopy (TCS SP8, Leica).

### Excitatory input stimulation at layer 5 pyramidal cell synapses

Excitatory postsynaptic currents (EPSCs) were evoked by extracellular stimulation in the vicinity of basal dendrites of layer 5 pyramidal cells (PCs) within layer 5 using a (ISO-Pulser ISOP1, AD-Elektronik, Buchenbach, Germany). Inputs were stimulated repetitively with a pattern consisting of a train of 30 pulses at 50 Hz followed by single pulses at different inter-stimulus intervals to probe recovery. Finally, 20 s after the 50 Hz train, a pair of stimuli with an inter-stimulus interval of 20 ms was delivered (Fig. 1B). This stimulus pattern (train + paired-pulses) was repeated every 45 s for ∼10 min before LTP induction (before LTP baseline period) and up to 25 minutes after LTP induction (after LTP baseline period), also referred to as early LTP or short-term potentiation (86, 87). To isolate EPSCs, the holding potential was set close to the Cl^−^-equilibrium potential (approximately −69 mV). Bath application of SR95531 induced “epileptic” activity during recordings (data not shown) and was therefore not applicable. In a subset of experiments, 20 µM SR95531 were locally pressure applied via the stimulation electrode to block local GABA_A_ receptors. LTP induced changes in EPSC size and short-term plasticity - were not significantly different between cells with and without local blockade of GABA_A_ receptors (Fig. S1) and data were therefore pooled. For comparisons before and after LTP induction, EPSCs during the last 9 min before induction (“before” LTP condition) and all EPSCs after induction (“after” LTP condition) were analyzed.

For paired recordings of synaptically connected layer 5 pyramidal cells (Fig. S3), presynaptic cells were stimulated by 1.5 ms long voltage steps in on-cell (200 – 400 mV) or whole-cell (100 mV) configuration as described previously (40, 82). Stimulation-triggered EPSCs were recorded in postsynaptic pyramidal cells in whole-cell configuration.

### LTP induction at layer 5 pyramidal cell synapses

To elicit electrically-induced LTP, pre- and post-synaptic activity was paired in current current-clamp mode and the bridge balance was set to 100%. The LTP induction protocol consisted of 30 pre-post pairings at 0.1 Hz (Fig. 1B). Presynaptic activity was elicited by extracellular stimulation of the inputs (8 pulses at 50 Hz) and was paired with a 2 ms delayed postsynaptic depolarization induced by a 200 ms long current injection into the postsynaptic cell to evoke AP firing (9, 10). The amplitude of the injected current was adjusted to achieve an AP firing frequency ≥35 Hz. Presynaptic stimulation and postsynaptic APs were not precisely timed. In between pairings, the membrane voltage (V_m_) of the postsynaptic cell was adjusted to approximately −70 mV.

### Excitatory input stimulation and plasticity induction at hippocampal CA1 pyramidal cells

After establishing a postsynaptic whole-cell voltage-clamp recording in visually identified CA1 pyramidal cells, EPSCs were evoked by extracellular stimulation of Schaffer collateral-commissural fibers in the stratum radiatum. The stimulation protocol and further experimental procedures were the same as for layer 5 pyramidal cells described above. In a subset of experiments, the CA3 region was cut to reduce recruitment of inhibitory inputs and the GABA_A_R antagonist SR95331 was bath applied through the perfusion system.

EPSCs potentiation was induced either by theta burst stimulation or by tetanic stimulation (200 Hz) of synaptic inputs while the postsynaptic CA1 neuron was held in current-clamp configuration. Theta burst stimulation included 4 trains, each containing 10 bursts of 4 stimuli at 100 Hz delivered every 200 ms. Trains were repeated every 10 s. The tetanic stimulation protocol consisted of 10 stimulation episodes, each containing 40 pulses at 200 Hz. The inter-episode interval was 5 s. In both cases, input stimulation was paired with a supra-threshold current injection into the postsynaptic CA1 neuron to evoke spiking activity (AP firing frequency ≥35 Hz). Presynaptic stimulation and postsynaptic APs were not precisely timed. In between pairings, the membrane voltage (V_m_) of the postsynaptic cell was adjusted to approximately −70 mV.

### Pharmacological induction of vesicular release potentiation

For pharmacological induction of EPSC potentiation, forskolin and PDBu were first dissolved in DMSO and further diluted in ACSF to a final concentration of 40 µM and 1 µM, respectively. After establishing the before-LTP-induction (baseline) period (see above), forskolin or PDBu containing ACSF were washed in through the perfusion system while continuing input stimulation. EPSCs recorded after 4.5 minutes of wash-in were used for analysis of the after induction period. To exclude possible effects of DMSO onto synaptic strength, control ACSF was supplemented with an identical amount of DMSO (≤0.1%) as forskolin-or PDBu-containing ACSF.

### Induction of homeostatic plasticity in neuronal cell culture

Homeostatic plasticity was induced by adding 2 µM Tetrodotoxin (TTX) to the culture medium for 48 hours prior to recordings. TTX was first diluted in 100 µL of fresh medium per treated well and subsequently added to the respective wells containing 1 mL of medium. For control condition, only 100 µL of fresh medium without TTX were added to the respective wells.

### Data analysis

#### EPSC analysis

EPSCs peak amplitudes were analyzed for each repetition with custom-written Igor Pro routines (WaveMetrics, Lake Oswego, OR, USA). To test for successful LTP induction, the first EPSCs in response to the paired-pulses and 30 pulses at 50 Hz were analyzed and their mean amplitudes before (9 min period before induction) and after LTP induction (all measured EPSCs after induction) were compared using a Student’s *t*-test. For the plots of first EPSC vs. recording time (Figs.1D, 5D and E and Fig. S3D), all EPSCs in a given cell were normalized to the corresponding average EPSC amplitude measured before LTP induction. Afterwards, the normalized EPSCs obtained within a period of 4.5 min (Fig.1D), 2.5 min (Fig. 5D and E) or 3 min (Fig. S3D) were averaged across all cells. PPR and steady-state amplitudes were calculated for each stimulation individually and averaged thereafter.

#### Recovery time constant

To evaluate the time constant of recovery from depression (τ recovery), EPSCs in response to high-frequency stimulation were analyzed for each repetition within the before and after plasticity induction period, averaged and normalized to the first EPSC amplitude of the high-frequency EPSC train. For each cell, the time course of recovery of the mean normalized EPSC amplitudes was fit with a mono- or biexponential function. A biexponential fit was performed only if (i) the ratio of the slow and fast time constant was larger than 3, (ii) χ^2^ of the bi-exponential fit was >4% better than χ^2^ of the mono-exponential fit and (iii) the amplitude fraction of each exponential component was larger than 0.05. In case of a bi-exponential fit, a weighted time constant (τ_wd_) was calculated as τ_wd_ = ((A_fast_ * τ_fast_) + (A_slow_ * τ_slow_)) / (A_fast_ + A_slow_), where τ_fast_ and τ_slow_ are the fast and slow time constants, and A_fast_ and A_slow_ are the amplitudes of the two components, respectively. In rare cases, exponential fits to the recovery time course yielded poor results and such cells were excluded for further analysis.

#### RRP size estimation

To evaluate the size of the RRP the Schneggenburger-Meyer-Neher methode (SMN plot) was used, which assumes EPSC depression due to RRP depletion and constant pool replenishment (35, 88). The total number of readily releasable vesicles was estimated for each cell from the y-intercept of extrapolated linear regressions to the last 5 cumulative steady-state EPSCs before and after LTP induction (Fig. 2F). Alternatively, individual RRP sizes were estimated by using the Elmqvist–Quastel method (EQ plot, 39), which assumes a constant fraction of vesicle pool consumption per AP and negligible vesicle pool replenishment during the early phase of the stimulus train. RRP sizes per cell were estimated from the x-intercept of the extrapolated linear regression fit to the first few EPSCs plotted as a function of the cumulative previous release (35, 89). Two cells could not be analyzed with the EQ method due to their nonlinear relationship between EPSC and cumulative EPSC during stimulus train onset and were excluded.

#### sEPSCs detection and analysis

Spontaneous EPSCs (sEPSCs) were recorded repeatedly for 5 s at different time points during the before and after plasticity induction period. sEPSCs were detected with the template-matching algorithm implemented in the NeuroMatic plug-in (90) (Version 3) for Igor Pro (WaveMetrics, Lake Oswego, OR, USA). Event number and amplitudes were measured in all repetitions and pooled per condition. The median sEPSC amplitude and frequency were calculated for each cell.

### Short-term plasticity models

#### Description of the models

The sequential model was implemented exactly as described previously in eqs. 5 and 42-46 of ref. (33) (cf. Fig. 3A). The parallel model was implemented according to the following equations similar to the previously described model 3 of ref. (37) and the “simple two-pool model” of ref. (91) (cf. Fig. S4A):

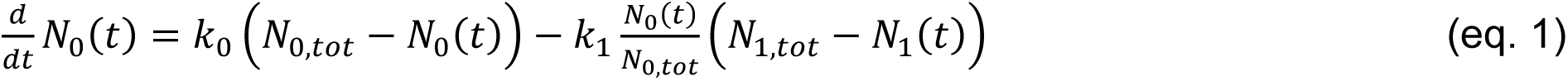

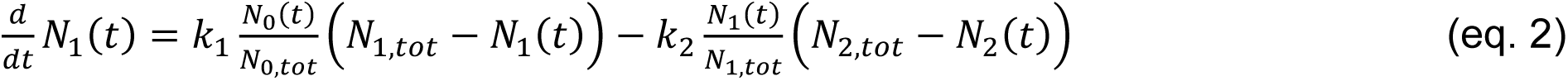

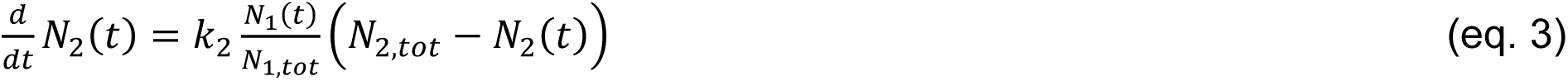

#### Parameter optimization for layer 5 pyramidal cell synapses

To optimize the free model parameters, we used a combination of manual adjustment and a simplex minimization algorithm while fitting the short-term plasticity model to EPSC amplitudes in response to stimulus trains at frequencies ranging from 0.5 to 50 Hz as well as in response to single stimuli delivered subsequently to conditioning stimulation at various recovery intervals (Figs. 3B and S4B). The quality of the fits was estimated by the sum of the squared differences between model fit and experimental data. Because of the mechanistic importance of the PPR and time course of recovery from depression, weighting factors were applied during the fit. In particular, the first two EPSCs were weighted with the factor of five and the EPSCs during the recovery from depression were weighted with a factor of three. Furthermore, because the subsequent analysis of the change of the parameters during LTP was restricted to 50 Hz trains and recovery following 50 Hz stimulation, each EPCS of the 50 Hz data was weighted with an additional factor of three.

The resulting best-fit parameters for the sequential model were: p_fusion_=0.5438, k_1,rest_=2.9653 s^-1^, b_1_=2.6177 s^-1^, k_2,rest_=0.3980 s^-1^, b_2_=1.0904 s^-1^, κ=0.1609 (the fraction of SP_LS_ transferred to SP_TSL_ per AP), b_3_ = 12.8963 s^-1^ (inverse of “decay τ of TSL”). For the Ca^2+^ dynamics, the parameters of the amplitude of “effective [Ca^2+^]” was a_Ca_=209.176 nM and the decay time constant of “effective [Ca^2+^]” was τ_Ca_=0.1395 s. The Ca^2+^-dependence of the ES→LS transition was defined according to eq. 42 of ref (33) with σ_1_=0.0732/(a_Ca_ τ_Ca_) and K_0.5_=612.018 nM. The Ca^2+^-dependence of the LS→TS transition was defined according to eq. 5 of ref. (33) with σ_2_=0.1051/(a_Ca_ τ_Ca_). The parameters for the change in p_fusion_ (eq. 37-41 in ref. (33)) were k_y_=71.4285 s^-1^, y_max_=1.32, y_inc,1_=0.39, and z _dec,1_=0. The resulting state occupancy of LS and TS vesicles was 0.44 and 0.16. For this initial parameter optimization based on normalized train data with various stimulation frequencies (Fig. 3B), the first EPSC amplitude was constrained to 1 by setting N_tot_ to

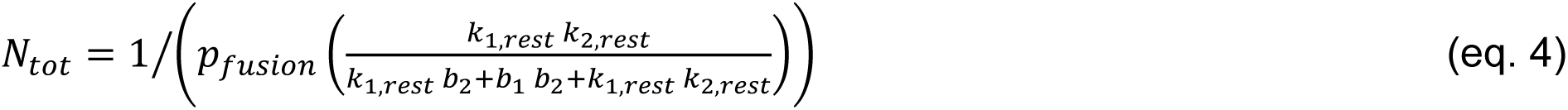

(i.e. N_tot_=11.32).

The resulting best-fit parameters for the parallel model were: p_2_=0.6772 and N_2_=0.6746. p_1_ was constrained to 0.5*p_2_ (i.e. p_1_=0.3386). For this initial parameter optimization based on normalized train data with various stimulation frequencies (Fig. S4B), the first EPSC amplitude was constrained to 1 by constraining N_1_ to obey the equation 1=N_2_*p_2_+N_1_*p_1_ (i.e. N_1_=1.604061). N_0_ was 6.7111. k_2,rest_, k_1,rest_, and k_0_ were 1.5374 s^-1^, 45.5642 s^-1^, and 2.6410 s^-1^, respectively. The model also contained a phenomenological description of facilitation as described previously (33), where each action potential increases both p_2_ and p_1_ by the amount p_x,intial_ (1-p_x_), with x=1 and 2. p_2_ and p_1_ then decay mono-exponentially back to p_x,initial_ with a time constant of 29.38334 ms.

#### Parameter optimization for before and after LTP induction

To estimate the change in the parameters following LTP induction, only five of the free parameters of both models were automatically optimized with a simplex minimization algorithm for the 50 Hz train and recovery data of each connection before and after LTP induction. Identical start values were used for all traces both before and after LTP induction. To quantify the quality of the fits, the sum of square differences of each amplitude was used and the first two EPSCs were weighted with the factor of five and the EPSCs in the recovery were weighted with a factor of three.

The free parameters for the sequential model were: k_1,rest_, b_1_, k_2,rest_, b_2_, and p_fusion_ (cf. Fig. 3E). The resulting occupancies of LS and TS states were calculated as

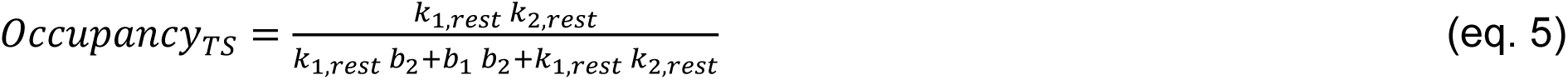

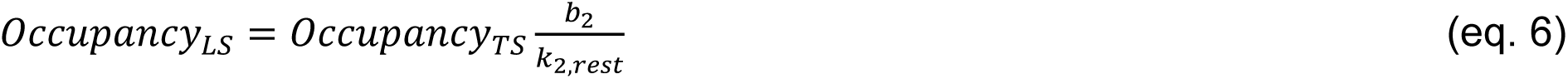

N_tot_ was constrained to eq. 4 multiplied by (a_before_ + a_after_)/2, where a_before_ and a_after_ are the median amplitude of the first EPSC of all responder cells before and after LTP, respectively. The free parameters for the parallel model were: N_2_, N_1_, k_2_, k_1_, and p_2_ (cf. Fig. S4E).

Changes in model parameters before and after LTP were presented as a relative difference (Figs. 3E and S4E) according to (eq. 7):

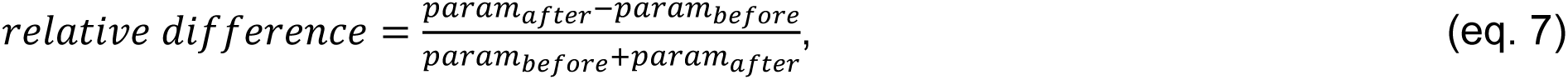

where param_before_ and param_after_ represent the corresponding parameters before and after LTP induction. This allows a comparison around 0 (no change) distributed symmetrically between -1 (decrease) and 1 (increase). In contrast, the ratio param_after_/param_before_ asymmetrically ranges from 0 to 1 (decrease) and from 1 to ∞ (increase) (92) (Figs. S5 and S8).

#### Linear un-mixing

As an alternative approach, we analyzed the contribution of each vesicles subpool to release during 50 Hz EPSC trains before and after LTP induction, as predicted by the models (Fig. S6). We therefore implemented a linear unmixing approach in analogy to (93). To calculate the release contributed by vesicles residing in specific pools at stimulus onset, we set each of the five free parameters to (param_before_ + param_after_)/2, where param_before_ and param_after_ are the median of the best-fit parameters of all responder cells before and after LTP induction, respectively. The rate “into” a given pool and the size of the downstream pool were set to zero. For example, to calculate the release component mediated by the pre-existing LS vesicles in the sequential model (cf. “pre-existing SV_LS_” in Fig. S6A-D), k_1_ and the initial occupancy of TS were set to zero. And, for example, to calculate the release component mediated by the pre-existing N_0_ vesicles in the parallel model (cf. “pre-existing SV_N0_” in Fig. S6E-H), k_0_, N_1_ and N_2_ were set to zero. For the parallel model, we in addition calculated the release component mediated “via the slots” of N_1_ and N_2_ vesicles by leaving all parameters unchanged but exclusively considering vesicles released from either the N_1_ or the N_2_ slots (Fig. S6I-L). Both alternative approaches for the parallel model revealed that the release component of the pre-existing N_2_ vesicles (Fig. S6E-H) and the release via the N_2_ “slot” (Fig. S6I-L) were significantly increased after LTP. The N_1_-release component was also increased but less so compared to the N_2_-release component. Because the N_1_-vesicles critically contribute to the steady-state release during the train, the slight elevation in the N_1_-release component is consistent with increased steady-state EPSC amplitude after LTP (cf. Fig. 2D). For the sequential model, induction of LTP increased the proportion of vesicles residing in TS at rest.

#### Correlations

To mechanistically analyze the correlations shown in Fig. 4A and C, we systematically varied the parameters of the models (Figs. 4E and S4F and G). We set each of the five free parameters to (param_before_ + param_after_)/2, where param_before_ and param_after_ are the median of the best-fit parameters of all responder cells before and after LTP, respectively. Next, each individual parameter was varied across a large range with hundred logarithmically spaced intervals. From these 100 release traces for each parameter, we automatically determined with Mathematica 12 (Wolfram Research, Champaign, IL, USA) the initial EPSC amplitude, PPR, steady-state EPSC, and time constant of recovery (by fitting mono- and biexponential functions and using the above-described criteria to choose τ_mono_ or τ_weighted_).

#### Implementation of the modelling

Both models were implemented in C++ using the compiler of XCode 14 on macOSX 13 (Apple Inc., Cupertino, CA, USA). The required computational time for the minimization of the five free parameters of the models for the EPSC amplitude before and after LTP induction (50 Hz train & recovery) was less than a minute for all 21 responder connections. The results of the minimization were visualized with Mathematica 12 (Wolfram Research, Champaign, IL, USA).

### Statistics

All pairwise comparisons between values before and after LTP induction were done by Wilcoxon signed rank tests. For unpaired comparisons between responder and non-responder cells Mann-Whitney rank sum tests were applied. For the multiple comparisons in Fig. 1G, Kruskal-Wallis one-way analysis of variance on ranks was performed followed by Dunn’s posthoc pairwise tests. All correlations of first EPSC size and PPR, and normalized steady-state EPSC and τ recovery (Figs. 4, 5 and S3) were tested for statistical significance using Spearman’s rank order correlation tests (r_s_ denotes the corresponding correlation coefficient). All values are presented as median [first quartile, third quartile] if not stated otherwise. Calculations of statistical tests were performed with jamovi (https://www.jamovi.org).

## Acknowledgements

This work was supported by the Deutsche Forschungsgemeinschaft (DFG, German Research Foundation; HA6386/10-1 to S.H.) and under Germany’s Excellence Strategy (EXC 2067/1-390729940) and by a European Research Council Consolidator Grant (ERC CoG 865634) to S.H.

## References

1. S. J. Martin, P. D. Grimwood, R. G. M. Morris, Synaptic Plasticity and Motor Learning. Neurosci. 23, 649–711 (2000).

2. H. R. Monday, T. J. Younts, P. E. Castillo, Long-Term Plasticity of Neurotransmitter Release: Emerging Mechanisms and Contributions to Brain Function and Disease. Annu Rev Neurosci 41, 299–322 (2018).

3. R. D. Hawkins, E. R. Kandel, S. A. Siegelbaum, Learning to modulate transmitter release: Themes and variations in synaptic plasticity. Annu. Rev. Neurosci. 16, 625–665 (1993).

4. P. E. Castillo, Presynaptic LTP and LTD of Excitatory and Inhibitory Synapses. Cold Spring Harb Perspect Biol 4, 5728–5729 (2012).

5. Y. Yang, N. Calakos, Presynaptic long-term plasticity. Front. Synaptic Neurosci. 5 (2013).

6. R. A. Nicoll, A Brief History of Long-Term Potentiation. Neuron 93, 281–290 (2017).

7. R. Enoki, Y. ling Hu, D. Hamilton, A. Fine, Expression of Long-Term Plasticity at Individual Synapses in Hippocampus Is Graded, Bidirectional, and Mainly Presynaptic: Optical Quantal Analysis. Neuron 62, 242–253 (2009).

8. T. V. P. Bliss, G. L. Collingridge, Expression of NMDA receptor-dependent LTP in the hippocampus: bridging the divide. Mol. Brain 6, 1–14 (2013).

9. H. Markram, M. Tsodyks, Redistribution of synaptic efficacy between neocortical pyramidal neurons. Nature 382, 807–810 (1996).

10. J. Sjöström, G. G. Turrigiano, S. B. Nelson, Multiple forms of long-term plasticity at unitary neocortical layer 5 synapses. Neuropharmacology 52, 176– 84 (2007).

11. M. Eder, W. Zieglgänsberger, H. U. Dodt, Neocortical long-term potentiation and long-term depression: Site of expression investigated by infrared-guided laser stimulation. J. Neurosci. 22, 7558–7568 (2002).

12. H. Markram, J. Lübke, M. Frotscher, B. Sakmann, Regulation of synaptic efficacy by coincidence of postsynaptic APs and EPSPs. Science (80-.). 275, 213–215 (1997).

13. B. M. Kampa, J. J. Letzkus, G. J. Stuart, Requirement of dendritic calcium spikes for induction of spike-timing-dependent synaptic plasticity. J. Physiol. 574, 283–290 (2006).

14. J. A. D’amour, R. C. Froemke, Inhibitory and excitatory spike-timing-dependent plasticity in the auditory cortex. Neuron 86, 514–528 (2015).

15. P. J. Sjöström, G. G. Turrigiano, S. B. Nelson, Rate, Timing, and Cooperativity Jointly Determine Cortical Synaptic Plasticity. Neuron 32, 1149–1164 (2001).

16. L. F. Abbott, S. B. Nelson, Synaptic plasticity: Taming the beast. Nat. Neurosci. 3, 1178–1183 (2000).

17. P. S. Kaeser, W. G. Regehr, The readily releasable pool of synaptic vesicles. Curr Opin Neurobiol. 43, 63–70 (2017).

18. A. Witkowska, L. P. Heinz, H. Grubmüller, R. Jahn, Tight docking of membranes before fusion represents a metastable state with unique properties. Nat. Commun. 12, 2–8 (2021).

19. A. Witkowska, S. Spindler, R. G. Mahmoodabadi, V. Sandoghdar, R. Jahn, Differential Diffusional Properties in Loose and Tight Docking Prior to Membrane Fusion. Biophys. J. 119, 2431–2439 (2020).

20. F. Michelassi, H. Liu, Z. Hu, J. S. Dittman, A C1-C2 Module in Munc13 Inhibits Calcium-Dependent Neurotransmitter Release. Neuron 95, 577–590.e5 (2017).

21. N. Lipstein, et al., Munc13-1 is a Ca2+-phospholipid-dependent vesicle priming hub that shapes synaptic short-term plasticity and enables sustained neurotransmission. Neuron 109, 1–21 (2021).

22. M. Camacho, et al., Control of neurotransmitter release by two distinct membrane-binding faces of the munc13-1 c1c2b region. Elife 10, 1–34 (2021).

23. B. Quade, et al., Membrane bridging by munc13-1 is crucial for neurotransmitter release. Elife 8, 1–30 (2019).

24. E. Neher, N. Brose, Dynamically Primed Synaptic Vesicle States: Key to Understand Synaptic Short-Term Plasticity. Neuron 100, 1283–1291 (2018).

25. M. Silva, V. Tran, A. Marty, Calcium-dependent docking of synaptic vesicles. Trends Neurosci. 44, 1–14 (2021).

26. S. Chang, T. Trimbuch, C. Rosenmund, Synaptotagmin-1 drives synchronous Ca 2+-triggered fusion by C 2 B-domain-mediated synaptic-vesicle-membrane attachment. Nat. Neurosci. 21, 33–42 (2018).

27. C. Imig, et al., The Morphological and Molecular Nature of Synaptic Vesicle Priming at Presynaptic Active Zones. Neuron 84, 416–431 (2014).

28. J. H. Jung, L. M. Kirk, J. N. Bourne, K. M. Harris, Shortened tethering filaments stabilize presynaptic vesicles in support of elevated release probability during LTP in rat hippocampus. Proc. Natl. Acad. Sci. U. S. A. 118, 1–8 (2021).

29. T. Miki, et al., Actin- and Myosin-Dependent Vesicle Loading of Presynaptic Docking Sites Prior to Exocytosis. Neuron 91, 808–823 (2016).

30. E. Hanse, B. Gustafsson, Vesicle release probability and pre-primed pool at glutamatergic synapses in area CA1 of the rat neonatal hippocampus. J. Physiol. 531, 481–493 (2001).

31. O. M. Schlüter, J. Basu, T. C. Südhof, C. Rosenmund, Rab3 superprimes synaptic vesicles for release: Implications for short-term synaptic plasticity. J. Neurosci. 26, 1239–1246 (2006).

32. H. Taschenberger, A. Woehler, E. Neher, Superpriming of synaptic vesicles as a common basis for intersynapse variability and modulation of synaptic strength. Proc. Natl. Acad. Sci. U. S. A. 113, E4548–E4557 (2016).

33. K.-H. Lin, H. Taschenberger, E. Neher, A sequential two-step priming scheme reproduces diversity in synaptic strength and short-term plasticity. Proc Natl Acad Sci U S A 119, (34) e2207987119 (2022).

34. S. Chanda, M. A. Xu-Friedman, A low-affinity antagonist reveals saturation and desensitization in mature synapses in the auditory brain stem. J. Neurophysiol. 103, 1915–1926 (2010).

35. E. Neher, Merits and Limitations of Vesicle Pool Models in View of Heterogeneous Populations of Synaptic Vesicles. Neuron 87, 1131–1142 (2015).

36. S. Hallermann, R. A. Silver, Sustaining rapid vesicular release at active zones: Potential roles for vesicle tethering. Trends Neurosci. 36, 185–194 (2013).

37. S. Hallermann, et al., Bassoon speeds vesicle reloading at a central excitatory synapse. Neuron 68, 710–723 (2010).

38. C. Keine, et al., Presynaptic Rac1 controls synaptic strength through the regulation of synaptic vesicle priming. Elife 11, e81505 (2022).

39. T. Branco, K. Staras, The probability of neurotransmitter release: variability and feedback control at single synapses. Nat. Rev. Neurosci. 2009 105 **10**, 373–383 (2009).

40. G. Bornschein, J. Eilers, H. Schmidt, Neocortical High Probability Release Sites Are Formed by Distinct Ca2+ Channel-to-Release Sensor Topographies during Development. Cell Rep. 28, 1410–1418.e4 (2019).

41. R. A. Nicoll, D. Schmitz, Synaptic plasticity at hippocampal mossy fibre synapses. Nat. Rev. Neurosci. 6, 863–876 (2005).

42. M. Orlando, et al., Recruitment of release sites underlies chemical presynaptic potentiation at hippocampal mossy fiber boutons. PLoS Biol. 19, 1–29 (2021).

43. J. T. R. Isaac, R. A. Nicoll, R. C. Malenka, Evidence for silent synapses: Implications for the expression of LTP. Neuron 15, 427–434 (1995).

44. D. Liao, R. H. Scannevin, R. Huganir, Activation of silent synapses by rapid activity-dependent synaptic recruitment of AMPA receptors. J. Neurosci. 21, 6008–6017 (2001).

45. S. Rey, V. Marra, C. Smith, K. Staras, Nanoscale Remodeling of Functional Synaptic Vesicle Pools in Hebbian Plasticity. Cell Rep. 30, 2006–2017.e3 (2020).

46. S. S. Zakharenko, L. Zablow, S. A. Siegelbaum, Visualization of changes in presynaptic function during long-term synaptic plasticity. Nat. Neurosci. 4, 711– 717 (2001).

47. J. del Castillo, B. Katz, Statistical factors involved in neuromuscular facilitation and depression. J Physiol. 24, 574–585 (1954).

48. C. Pulido, A. Marty, Quantal fluctuations in central mammalian synapses: Functional role of vesicular docking sites. Physiol. Rev. 97, 1403–1430 (2017).

49. C. Imig, et al., Ultrastructural Imaging of Activity-Dependent Synaptic Membrane-Trafficking Events in Cultured Brain Slices. Neuron 108, 843–860.e8 (2020).

50. D. Vandael, C. Borges-Merjane, X. Zhang, P. Jonas, Short-Term Plasticity at Hippocampal Mossy Fiber Synapses Is Induced by Natural Activity Patterns and Associated with Vesicle Pool Engram Formation. Neuron 107, 509–521.e7 (2020).

51. V. N. Murthy, T. Schikorski, C. F. Stevens, Y. Zhu, Inactivity produces increases in neurotransmitter release and synapse size. Neuron 32, 673–682 (2001).

52. V. N. Murthy, T. J. Sejnowski, C. F. Stevens, Heterogeneous release properties of visualized individual hippocampal synapses. Neuron 18, 599–612 (1997).

53. T. Sakaba, E. Neher, Calmodulin mediates rapid recruitment of fast-releasing synaptic vesicles at a calyx-type synapse. Neuron 32, 1119–1131 (2001).

54. J. S. Lee, W. K. Ho, E. Neher, S. H. Lee, Superpriming of synaptic vesicles after their recruitment to the readily releasable pool. Proc. Natl. Acad. Sci. U. S. A. 110, 15079–15084 (2013).

55. A. Ritzau-Jost, et al., Ultrafast action potentials mediate kilohertz signaling at a central synapse. Neuron 84, 152–163 (2014).

56. S. Hallermann, M. Heckmann, R. J. Kittel, Mechanisms of short-term plasticity at neuromuscular active zones of Drosophila. HFSP J. 4, 72–84 (2010).

57. B. Pan, R. S. Zucker, A General Model of Synaptic Transmission and Short-Term Plasticity. Neuron 62, 539–554 (2009).

58. R. Fernández-Busnadiego, et al., Quantitative analysis of the native presynaptic cytomatrix by cryoelectron tomography. J. Cell Biol. 188, 145–156 (2010).

59. R. P. Costa, B. E. P. Mizusaki, P. J. Sjöström, M. C. W. van Rossum, Functional consequences of pre- and postsynaptic expression of synaptic plasticity. Philos. Trans. R. Soc. B Biol. Sci. 372 (2017).

60. J. S. Rothman, L. Cathala, V. Steuber, R. A. Silver, Synaptic depression enables neuronal gain control. Nature 457, 1015–1018 (2009).

61. R. Fukaya, et al., Increased vesicle fusion competence underlies long-term potentiation at hippocampal mossy fiber synapses. 2, 1–14 (2023).

62. A. Betz, et al., Munc13-1 is a presynaptic phorbol ester receptor that enhances neurotransmitter release. Neuron 21, 123–136 (1998).

63. J. Basu, A. Betz, N. Brose, C. Rosenmund, Munc13-1 C1 domain activation lowers the energy barrier for synaptic vesicle fusion. J. Neurosci. 27, 1200– 1210 (2007).

64. J.-S. Rhee, et al., “Phorbol Ester-and Diacylglycerol-Induced Augmentation of Transmitter Release Is Mediated by Munc13s and Not by PKCs regions and involve successive, spatially segregated modulations of pre-and postsynaptic processes (Ma-lenka and Nicoll, 1999). Presynaptic” (2002).

65. J. S. Dittman, Unc13: a multifunctional synaptic marvel. Curr. Opin. Neurobiol. 57, 17–25 (2019).

66. L. Deng, P. S. Kaeser, W. Xu, T. C. Südhof, RIM proteins activate vesicle priming by reversing autoinhibitory homodimerization of munc13. Neuron 69, 317–331 (2011).

67. A. Betz, et al., Functional interaction of the active zone proteins Munc13-1 and RIM1 in synaptic vesicle priming. Neuron 30, 183–196 (2001).

68. J. Lu, et al., Structural basis for a Munc13-1 homodimer to Munc13-1/RIM heterodimer switch. PLoS Biol. 4, 1159–1172 (2006).

69. M. Camacho, et al., Heterodimerization of Munc13 C2A domain with RIM regulates synaptic vesicle docking and priming. Nat. Commun. 8, 1–13 (2017).

70. E. Fourcaudot, et al., cAMP/PKA signaling and RIM1α mediate presynaptic LTP in the lateral amygdala. Proc. Natl. Acad. Sci. U. S. A. 105, 15130–15135 (2008).

71. P. E. Castillo, S. Schoch, F. Schmitz, T. C. Südhof, R. C. Malenka, RIM1α is required for presynaptic long-term potentiation. Nature 415, 327–330 (2002).

72. J. A. Müller, et al., A presynaptic phosphosignaling hub for lasting homeostatic plasticity. Cell Rep. 39, 110696 (2022).

73. 73. P. S. Kaeser, et al., “RIM1 phosphorylation at serine-413 by protein kinase A is not required for presynaptic long-term plasticity or learning” (2008).

74. P. E. Castillo, et al., Rab3A is essential for mossy fibre long-term potentiation in the hippocampus. Nature 388, 590–593 (1997).

75. Y. Y. Huang, et al., Genetic evidence for a protein-kinase-A-mediated presynaptic component in NMDA-receptor-dependent forms of long-term synaptic potentiation. Proc. Natl. Acad. Sci. U. S. A. 102, 9365–9370 (2005).

76. M. Midorikawa, T. Sakaba, Kinetics of Releasable Synaptic Vesicles and Their Plastic Changes at Hippocampal Mossy Fiber Synapses. Neuron 96, 1033–1040.e3 (2017).

77. R. Fukaya, M. Maglione, S. J. Sigrist, T. Sakaba, Rapid Ca2+ channel accumulation contributes to cAMP-mediated increase in transmission at hippocampal mossy fiber synapses. Proc. Natl. Acad. Sci. U. S. A. 118, 1–11 (2021).

78. G. Malagon, T. Miki, V. Tran, L. C. Gomez, A. Marty, Incomplete vesicular docking limits synaptic strength under high release probability conditions. Elife 9, 1–18 (2020).

79. E. He, et al., Munc13-1 and Munc18-1 together prevent NSF-dependent de-priming of synaptic vesicles. Nat. Commun. 8, 1–10 (2017).

80. A. Eshra, H. Schmidt, J. Eilers, S. Hallermann, Calcium dependence of neurotransmitter release at a high fidelity synapse. Elife 10, 1–34 (2021).

81. N. Hosoi, T. Sakaba, E. Neher, Quantitative analysis of calcium-dependent vesicle recruitment and its functional role at the calyx of held synapse. J. Neurosci. 27, 14286–14298 (2007).

82. G. Bornschein, S. Brachtendorf, H. Schmidt, Developmental Increase of Neocortical Presynaptic Efficacy via Maturation of Vesicle Replenishment. Front. Synaptic Neurosci. 11, 1–10 (2020).

83. J. Bischofberger, D. Engel, L. Li, J. R. P. Geiger, P. Jonas, Patch-clamp recording from mossy fiber terminals in hippocampal slices. Nat. Protoc. 1, 2075–2081 (2006).

84. J. T. Ting, et al., Preparation of Acute Brain Slices Using an Optimized N-Methyl-D-glucamine Protective Recovery Method. J. Vis. Exp. 2018, 1–13 (2018).

85. A. Ritzau-Jost, et al., Large, stable spikes exhibit differential broadening in excitatory and inhibitory neocortical boutons. Cell Rep. 34, 108612 (2021).

86. K. M. Harris, Synaptic odyssey. J. Neurosci. 40, 61–80 (2020).

87. Uwe Frey, Richard G. M. Morris, Synaptic tagging and long-term potentiation. Nature 385, 533–536 (1997).

88. R. Schneggenburger, A. C. Meyer, E. Neher, Released fraction and total size of a pool of immediately available transmitter quanta at a calyx synapse. Neuron 23, 399–409 (1999).

89. D. Elmqvist, D. M. Quastel, A quantitative study of end-plate potentials in isolated human muscle. test. J. Physiol. 178, 505–529 (1965).

90. J. S. Rothman, R. A. Silver, Neuromatic: An integrated open-source software toolkit for acquisition, analysis and simulation of electrophysiological data. Front. Neuroinform. 12, 1–21 (2018).

91. A. Ritzau-Jost, et al., Apparent calcium dependence of vesicle recruitment. J. Physiol. 596, 4693–4707 (2018).

92. A. Schmid, et al., Activity-dependent site-specific changes of glutamate receptor composition in vivo. Nat. Neurosci. 11, 659–666 (2008).

93. E. Neher, H. Taschenberger, Non-negative Matrix Factorization as a Tool to Distinguish Between Synaptic Vesicles in Different Functional States. Neuroscience 458, 182–202 (2021).

