## supplemental data for "Fully-primed slowly-recovering vesicles mediate presynaptic LTP at neocortical neurons"

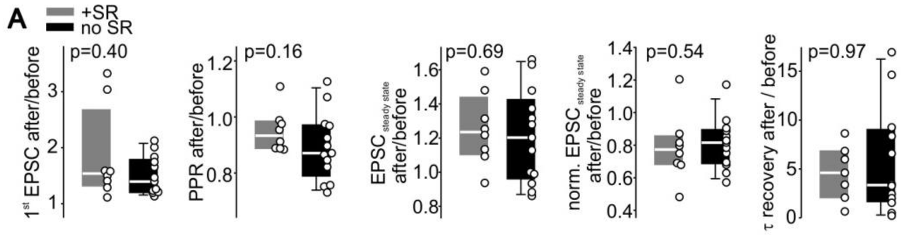


**Fig. S1**

(A) Boxplots and scatter dot plots of EPSC and short-term plasticity properties with (+SR) and without (no SR) local application of the $GABA_AR$ antagonist SR95531.

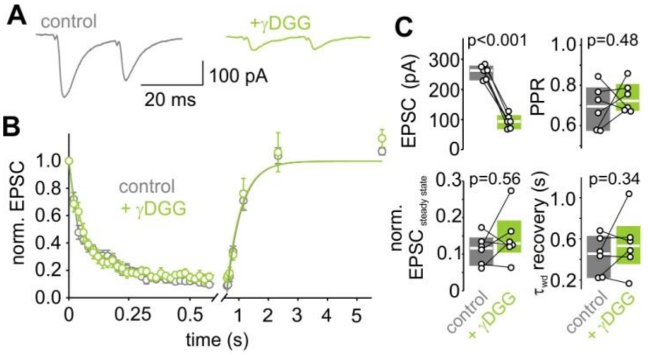


**Fig. S2 Postsynaptic saturation and desensitization do not contribute to short-term plasticity**

(A) Example average EPSC traces of paired-pulses before and after application of the low-affinity competitive AMPAR antagonist γDGG.

(B) Mean normalized 50 Hz EPSC trains and single EPSCs recorded at different recovery intervals were unchanged after γDGG application.

(C) Boxplots and individual EPSC amplitudes, short-term plasticity parameters and recovery time constants before (control) and after application of γDGG.

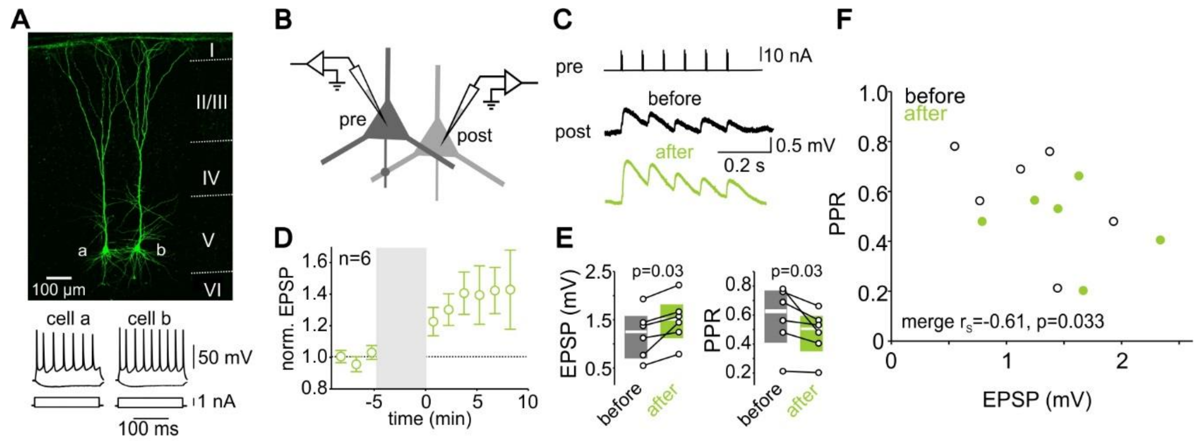


**Fig. S3 Electrical-induced LTP in two connected layer 5 pyramidal cells revealed by paired-recordings**

(A) Post hoc labeling of two biocytin-filled neocortical layer 5 pyramidal cells in rat cortical slices with the typical regular AP firing during current injection (bottom).

(B) Schematic of paired-recording configuration. Presynaptic cells were stimulated in on-cell or whole-cell configuration and corresponding EPSPs were recorded in the postsynaptic cell (for details see Methods).

(C) Example average traces of EPSPs in a layer 5 PC elicited by voltage stimulation of the presynaptic pyramidal cell in on-cell configuration before and after LTP induction.

(D) Normalized mean EPSP amplitudes of responder cells before and after LTP induction.

(E) Individual and median EPSP amplitudes and PPRs of responder cells before and after LTP induction.

(F) Scatter plots of PPR vs. initial EPSP amplitude in paired whole-cell recordings of two synaptically coupled layer 5 pyramidal cells before and after application of different LTP induction protocols.

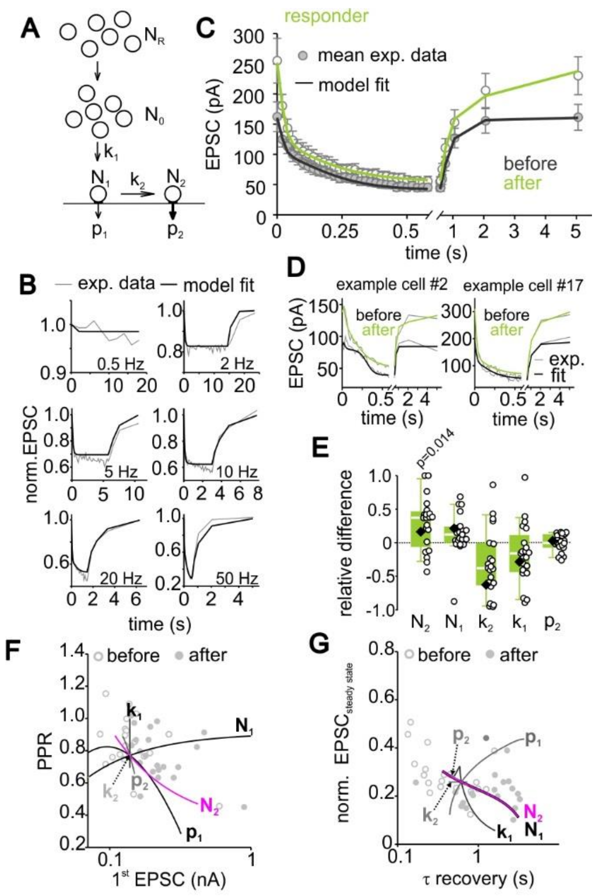

**Fig. S4 A parallel priming model indicates an increased number of superprimed vesicles following LTP induction**

(A) Parallel model containing 2 fusion-competent vesicle pools with normally primed vesicles ($N_1$) exhibiting a low release probability ($p_1$) and superprimed vesicles ($N_2$) with a high release probability ($p_2$), a supply pool ($N_0$) and infinite reserve pool ($N_R$). $N_1$ and $N_2$ refilling determined by the rate constants $k_1$ and $k_2$, respectively.

(B) Model fit to low frequency transmission.

(C) Model fit to the mean 50-Hz-train- and recovery EPSCs of responder cells before and after LTP.

(D) Model fits to individual 50-Hz-train- and recovery EPSCs before and after LTP in two example cells.

(E) LTP induced individual (circles) and median relative difference (eq.7) of the predicted model parameters (diamond = parameter obtained from model fit to the average of all responder cells). P values were calculated by comparison of respective parameters before and after LTP induction by Wilcoxon signed rank tests.

(F, G) Model predictions of the EPSC vs. PPR (F) and normalized steady-state EPSC vs. recovery (G) correlation by individually varying the model parameters.

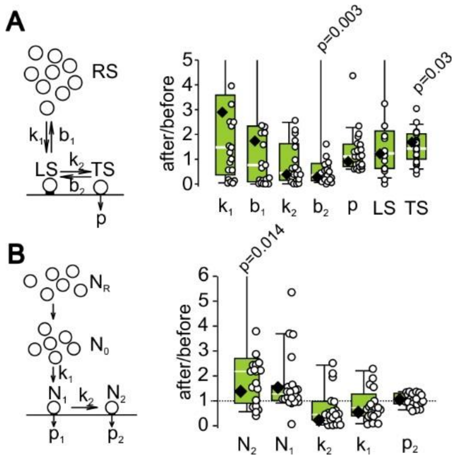


**Fig. S5 Ratio of LTP-induced changes of model parameter for comparison with relative difference in responder cells**

(A) Schematic of the sequential priming model (cf. Fig. 3, left) and LTP induced individual (circles) and median relative changes of the predicted model parameters (right; diamond = parameter obtained from model fit to the average of all responder cells).

(B) Schematic of the parallel priming model (cf. Fig. S4, left) and LTP induced individual (circles) and median relative changes of the predicted parallel model parameters (right; diamond = parameter obtained from model fit to the average of all responder cells)

P values were calculated by comparison of respective parameters before and after LTP induction by Wilcoxon signed rank tests.

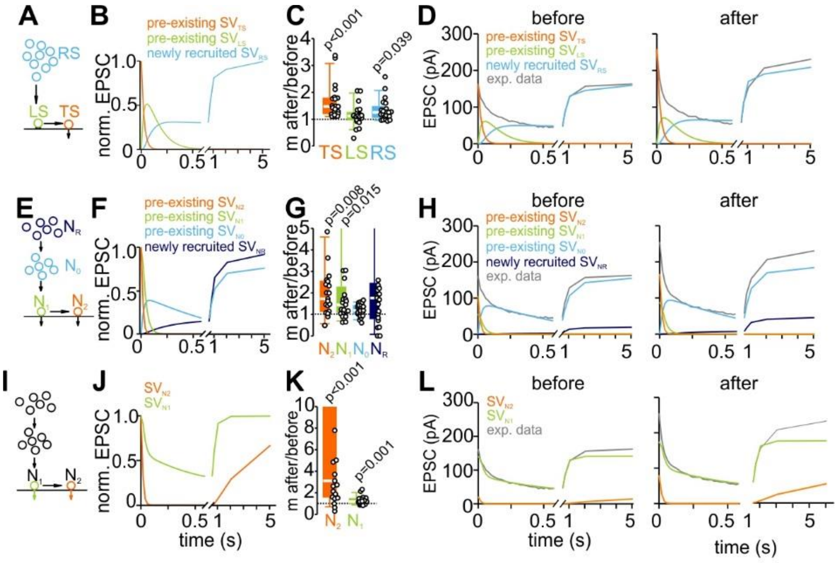

**Fig. S6 Linear unmixing-approach indicates a predominant increase of pre-existing fully-primed vesicles upon LTP**

(A, B) Schematic of the sequential model and predicted relative release from the individual pools during 50 Hz transmission and subsequent recovery of synaptic depression.

(C) Relative changes of the release from the respective subpools of the sequential model following LTP induction.

(D) Predicted contributions to release from the subpools before and after LTP induction in comparison to experimentally obtained total release.

(E, F) Schematic of the parallel model and predicted relative release from the individual pools during 50 Hz transmission and subsequent recovery of synaptic depression.

(G) Relative LTP-induced changes in contribution to release by the respective subpools for the parallel model.

(H) Predicted contributions to release by the subpools before and after LTP induction in comparison to experimentally obtained total release.

(I, J) Parallel model with only fusion-competent vesicles and predicted relative release from the two pools during 50 Hz transmission and subsequent recovery of synaptic depression.

(K) Relative LTP-induced changes in contribution to release by the respective subpools for the parallel model.

(L) Predicted contributions to release by the subpools before and after LTP induction in comparison to experimentally obtained total release.

P values were calculated by comparison of respective parameters before and after LTP induction by Wilcoxon signed rank tests.

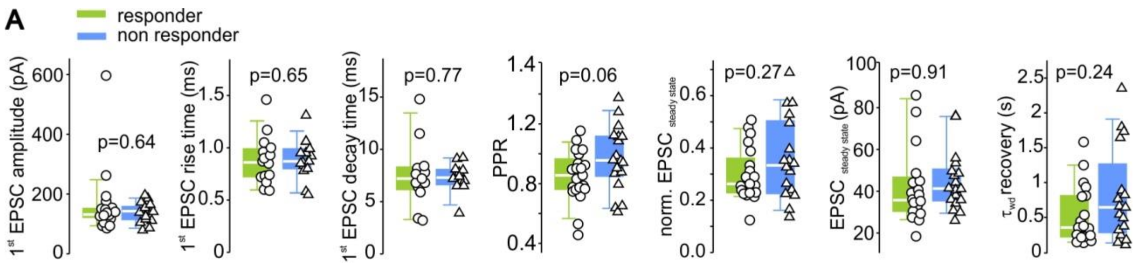


**Fig. S7 Properties of EPSCs and short-term plasticity of LTP responder and non-responder cells**

(A) Boxplots and individual data points of EPSC and short-term plasticity properties of responder (n=21) and non-responder (n=18) cells.

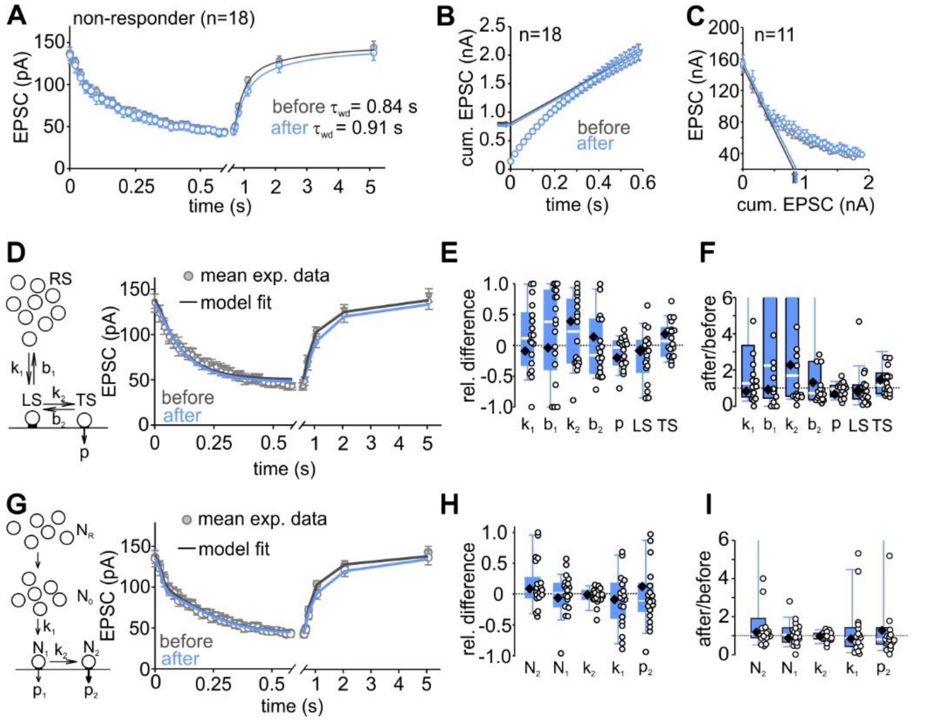


**Fig. S8 EPSC train properties of non-responder cells were unchanged during LTP**

(A) Mean normalized 50-Hz EPSC train and recovery EPSCs before and after the LTP induction protocol was applied.

(B, C) Estimates for RRP size obtained by the *SNM method* (B) and EQ (C) method were similar before and after LTP induction in non-responder cells.

(D) Schematic of the sequential priming and fusion model (left) and model fit to the average of high-frequency EPSC trains before and after LTP induction for non-responder cells (right).

(E, F) Individual and median relative difference (eq.7, E) and ratio (F) of the predicted sequential model parameters before and after the LTP induction protocol (diamond = parameter obtained from model fit to the average of all responder cells).

(G) Schematic of the parallel priming and fusion model (left) and model fit to the average of high-frequency EPSC trains before and after LTP induction for non-responder cells (right).

(H, I) Individual and median relative difference (eq.7, H) and ratio (I) of the predicted sequential model parameters before and after the LTP induction protocol (diamond = parameter obtained from model fit to the average of all responder cells).
